# Mutant huntingtin exon-1 impaired GTPCH and DHFR expression in plants and mice

**DOI:** 10.1101/2022.05.18.492514

**Authors:** Chiu-Yueh Hung, Chuanshu Zhu, Farooqahmed S. Kittur, Maotao He, Erland Arning, Jianhui Zhang, Asia J. Johnson, Gurpreet S. Jawa, Michelle D. Thomas, Tomas T. Ding, Jiahua Xie

**Affiliations:** Department of Pharmaceutical Sciences, Biomanufacturing Research Institute & Technology Enterprise, North Carolina Central University, Durham, NC 27707, USA.; College of Plant Protection, Northwest A&F University, Yangling, Shaanxi 712100, China.; Department of Pathology, Weifang Medical University, Weifang 261000, China.; Baylor Scott & White Research Institute, Institute of Metabolic Disease, Dallas, TX 75204, USA; DePuy Synthes Companies of Johnson & Johnson, West Chester, PA 19380, USA; Present address: University of North Carolina, Eshelman School of Pharmacy, Chapel Hill, NC 27599, USA

**Author notes:** These authors contributed equally to this work (Chiu-Yueh Hung, Chuanshu Zhu, Farooqahmed S. Kittur, Maotao He, Erland Arning).

**Keywords:** Dihydrofolate reductase, GTP cyclohydrolase I, Huntington’s disease, one-carbon metabolism, plant-based model, tetrahydrobiopterin biosynthesis.

## Abstract

Pathophysiology associated with Huntington’s disease (HD) has been studied extensively in various cell and animal models since the 1993 discovery of the mutant huntingtin (mHtt) with abnormally expanded polyglutamine (polyQ) tracts as the causative factor. However, the sequence of early pathophysiological events leading to HD still remains elusive. To gain new insights into the polyQ-induced early pathogenic events, we expressed Htt exon1 (Htt_ex1_) with a normal (21), or an extended (42 or 63) number of polyQ in tobacco plants, which lack an Htt ortholog to avoid any associated effects from endogenous *Htt*. Here, we show that transgenic plants accumulated Htt_ex1_ proteins with corresponding polyQ tracts, and that mHtt_ex1_ induced protein aggregation and affected plant growth, especially root and root hair development, in a polyQ length-dependent manner. Quantitative proteomic analysis of young roots from severely affected Htt_ex1_Q63 and unaffected Htt_ex1_Q21 plants showed that the most impaired protein by polyQ63 is a GTP cyclohydrolase I (GTPCH) along with many its related one-carbon (C_1_) metabolic pathway enzymes. GTPCH is a key enzyme involved in folate biosynthesis in plants and tetrahydrobiopterin (BH_4_) biosynthesis in mammals. Validating studies in 4-week-old R6/2 HD mice expressing a mHtt_ex1_ showed reduced levels of GTPCH and dihydrofolate reductase (DHFR, a key folate utilization/alternate BH_4_ biosynthesis enzyme), and impaired C_1_ and BH_4_ metabolisms. Our findings from mHtt_ex1_ plants and mice reveal impaired expressions of GTPCH and DHFR and contribute to a better understanding of mHtt-altered C_1_ metabolism and C_1_ interconnected BH_4_ metabolism leading to the pathogenesis of HD.

## Introduction

Huntington’s disease (HD) and eight other polyglutamine (polyQ)-mediated neurodegenerative diseases (NDDs) share the common features of pathological protein aggregation, and age- dependent progressive neurodegeneration^1, 2^. After years of intensive research following the discovery of abnormally expanded polyQ (>36Q) in huntingtin exon 1 (Htt_ex1_) being responsible for HD^3^, the underlying pathophysiological mechanisms are still not fully understood and no effective cure is available^4–6^. Limited study to unravel early cellular events affected by polyQ repeats may be one of key obstacles to understand the initiation and progression of the disease. Exploring novel systems might facilitate our understanding of disease initiation and progression processes.

Plants could be useful for studying polyQ repeat-induced pathology because both plant and animal cells are eukaryotic and share many similarities while plants have some uniquenesses.

Most importantly, plants naturally lack Htt homologs, and transgenic plants expressing Htt or mHtt would avoid any endogenous Htt’s effects, which will simplify the interpretation of polyQ effects compared to any HD animal models. Furthermore, recent molecular, biochemical and physiological studies have shown that plant root apices possess sensing functions, such as receiving, integrating and responding to signals from their environment, and that root hairs share several morphological and biochemical features with neurons, such as a single tubular-shaped cell, long extension tip-growth, high energy demand, cell polarity, action potential, and environmental sensing capacity^7, 8^. Therefore, roots and root hairs might be more susceptible to polyQ-induced toxicity than other organs/cells to exhibit phenotypic changes. In addition, plants also possess numerous experimental advantages, such as less ethical requirements, easy cell culture and transformation, and simple growth and seed maintenance. Plants have been utilized to study human diseases^9, 10^.

To explore the use of transgenic plants as a system to investigate the toxicity of abnormal polyQ repeats and early affected cellular processes, we first proved that expressing human *Htt_ex1_* with different “CAG” repeats in transgenic tobacco (*Nicotiana tabacum*) plants could accumulate Htt_ex1_ with the corresponding polyQ tracts. We then discovered that the toxicity of the expanded polyQ repeats to plant cells was dependent on the repeat length. The most intriguing observation was that polyQ63 inhibited root growth and root hair outgrowth while polyQ21 did not display such inhibitory effects. Quantitative proteomic analysis of young adventitious roots induced from shoots of Htt_ex1_Q63 and Htt_ex1_Q21 revealed that many differentially abundant proteins (DAPs) were enriched in various cellular pathways that have previously been implicated in HD models. Importantly, we discovered that GTP cyclohydrolase I (GTPCH), the rate-limiting enzyme catalyzing first step in the de novo folate biosynthesis, was the most affected protein by polyQ63. Moreover, we also observed many other DAPs that are involved in activating and transferring one carbon (C_1_) units to the various methylation pathways^11, 12^. Our findings of polyQ63 impaired GTPCH and its associated C_1_ metabolism are novel because previous studies have suggested a possible link between NDDs, including HD, and the disorder of folate-mediated C_1_ metabolism^13–,16^, but the molecular link connecting C_1_ metabolism to NDDs has been missing. In animals, GTPCH catalyzes the first and limiting step in the tetrahydrobiopterin (BH_4_) biosynthesis, and the latter servers as a co-factor for the production of the monoamine neurotransmitters^17^. Therefore, we set out to validate our novel findings from Htt_ex1_Q63 transgenic plants in the well-established R6/2 HD mice. We discovered that mHtt_ex1_ truly suppressed the levels of GTPCH as well as dihydrofolate reductase (DHFR), the key enzyme of folate utilization and alternate BH_4_ biosynthesis^15, 17^ with disturbed C_1_ and BH_4_ metabolisms in 4-week old R6/2 mice.

## Results

### Generation of *mHtt_ex1_* transgenic plants

*Htt_ex1_* with expanded “CAG” repeats that have been confirmed to be toxic and cause protein aggregation is commonly used to create HD models^18, 19^. We therefore expressed human *Htt_ex1_* containing normal (20), or abnormal (40 or 60) “CAG” repeats with additional 1, 2 or 3 “CAA” (also encoding Q) repeats, and a bacterial *uidA* gene encoding *β*-glucuronidase (GUS) (as a vector control) (Fig. 1A) in tobacco plants to produce transgenic plants named Htt_ex1_Q21, Htt_ex1_Q42, Htt_ex1_Q63, and GUS, respectively. PolyQ63 dramatically affected the number of shoots per explant and leaf color, and hampered plant growth and development, but did not affect the tissue culture response nor the sizes of produced pods and seeds (Fig. S1 and S2). PloyQ42 had less negative effects, while polyQ21 displayed no negative effects compared to GUS. Transgenes were confirmed to be stably integrated into the tobacco genome by PCR and properly transcribed by RT-PCR analysis (Fig. 1B and 1C; Fig. S3). Immunoblotting analysis of two PCR confirmed transgenic lines from each construct showed immunoreactive bands of sizes ∼20, ∼26 and ∼32 kD (Fig. 1D), twice their predicted sizes as also observed in a mouse HD model expressing *N*- terminal mHtt^20^. The authors^20^ suggested that the Htt peptides with polyQ easily form di- and trimers. The obtained results indicate that these stable transgenic plants accumulated Htt peptides, each with the expected polyQ length, and that the toxic effects of polyQ tracts on plant growth are length-dependent.

**Fig. 1.**
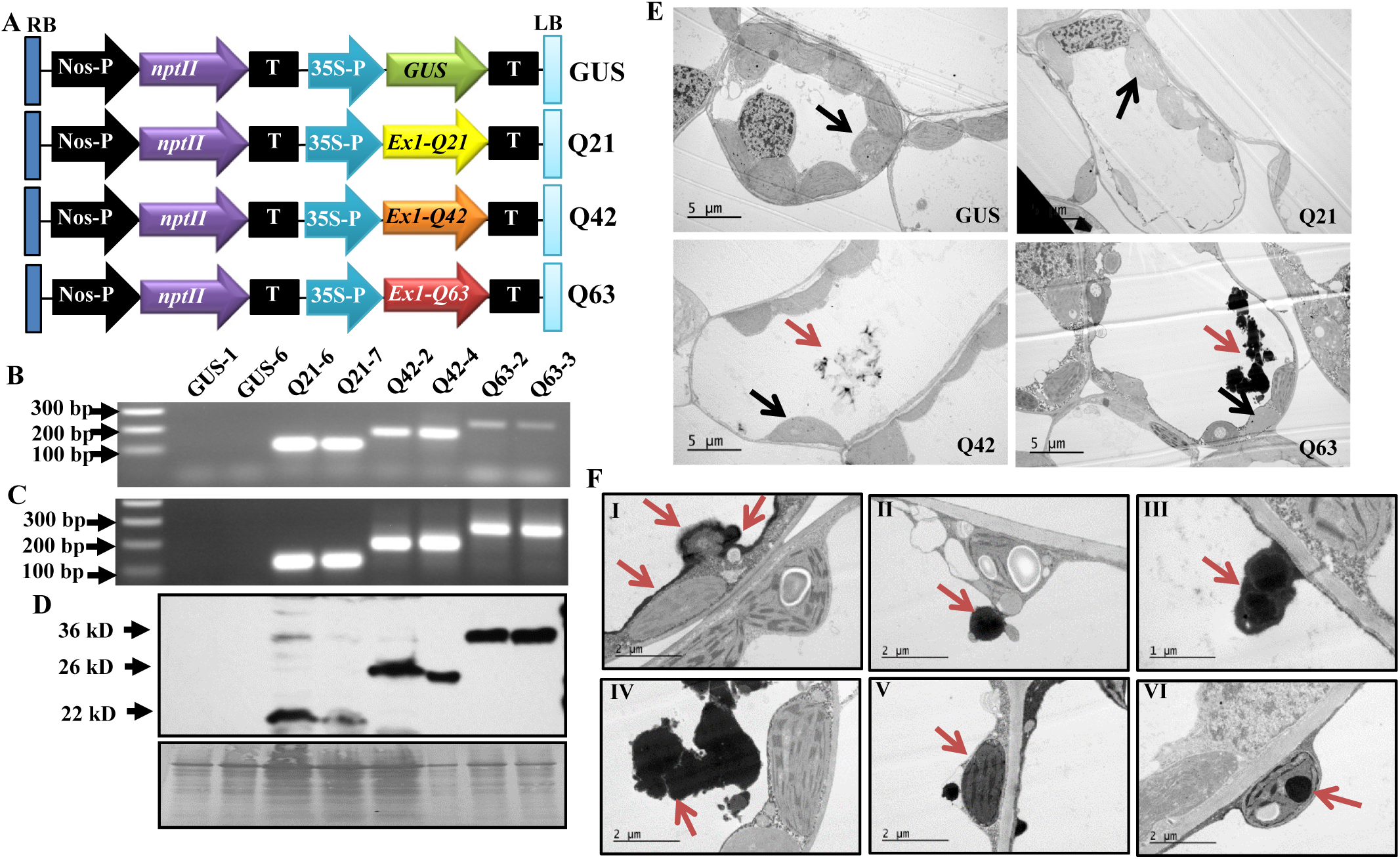
Genetic cassettes and transgenic plant characterization. **(A)** Schematic representation of four genetic cassettes used to transform tobacco plants. **(B-D)** Representative results of two transgenic lines per genetic cassette analyzed by genomic PCR (B), RT-PCR (C), and immunoblotting with anti-Htt antibody mEM48 (D). **(E)** Microstructures of transgenic leaf cells under electron microscope. Red arrows indicate protein aggregates while black arrows indicate chloroplasts. **(F)** Observation of different stages of aggregate formation in young leaves of Htt_ex1_Q63 plants under TEM: initiated around chloroplasts and along the tonoplast (I), formed aggregates (II, III) released into the vacuoles (IV). They are also observed in cytoplasm (V) and inside of chloroplasts (VI). Q21: Htt_ex1_Q21; Q42: Htt_ex1_Q42; and Q63: Htt_ex1_Q63.

### PolyQ63 and polyQ42 cause protein aggregation

The presence of neuronal aggregates/inclusion bodies is a pathological hallmark in HD patients and models^19, 21, 22^. To determine whether overexpressing mHtt_ex1_ could also cause protein aggregation, transgenic leaf cells were first used to observe aggregates. Aggregates were frequently observed in Htt_ex1_Q63 young leaf cells while only small aggregates were occasionally found in a few Htt_ex1_Q42 cells, whereas no aggregates were detected in either Htt_ex1_Q21 or GUS cells (Fig. 1E). In Htt_ex1_Q63, the aggregates were often localized in vacuoles, which appeared to be initiated around chloroplasts and along the tonoplasts with a progressive increase in size and then released into vacuoles (Fig. 1F). They were also observed in the cytoplasm and inside of the chloroplasts (Fig. 1F). We also noticed that Htt_ex1_Q63 had much smaller chloroplasts with roughly half the size observed in the GUS control (Fig. 1E).

Since mHtt_ex1_, ubiquitin and Hsp70 were reported to be present inside polyQ-aggregates in mammalian cell and animal models of HD^22, 23^, we investigated whether they are also present inside of the plant polyQ-aggregates by employing Blue Native-PAGE (BN-PAGE) immunoblotting and filter retardation assays. Both assays showed that mHtt_ex1_, ubiquitin and Hsp70 were present in Htt_ex1_Q63 lines (Fig. S4), consistent with the observations made in mammalian cell and animal models^22, 23^. The observed length-dependent polyQ-aggregates and detected mHtt_ex1_, ubiquitin and Hsp70 inside aggregates indicate that the plant-based protein aggregation model recapitulates a major pathogenic process of HD reported in animal models^18, 19^ and could be useful for dissecting the underlying mechanism of mHtt-induced toxicity.

### PolyQ63 affects root and root hair growth

Root hairs and neurons have been reported to share similar morphological and biochemical features^7, 8^, and both the axon extension and root hair tip-growth processes are tightly associated with tubular endoplasmic reticulum (ER) remodeling in the direction of cell elongation^8, 24–26^.

During the subculture and propagation of transgenic plants, we paid special attention to root growth. We did observe slow growth of induced adventitious roots from Htt_ex1_Q63 shoots with no root hairs in the maintenance medium (Fig. 2A and 2B). These phenomena prompted us to systematically investigate the effects of polyQ-repeat length on adventitious root emergence and subsequent growth. We found that the percentages of shoots with emerging roots among the four types of transgenic lines were not different (Fig. 2C). However, emerging roots from Htt_ex1_Q63 were shorter (∼0.5 cm) and devoid of root hairs on day 11 of the subculture, whereas those from Htt_ex1_Q42 had the same length (∼3 cm) as Htt_ex1_Q21 and GUS (Fig. 2D) but lesser root hair density (Fig. 2B). When the root hair emerging site from the root tip was measured, 43 out of 50 Htt_ex1_Q63 roots had no root hairs while all Htt_ex1_Q42, Htt_ex1_Q21, and GUS roots had root hairs with distances of 2.2-2.6 cm (Fig. 2E). When root tips were observed in detail, most of the Htt_ex1_Q63 root tips were found to be loosely organized while Htt_ex1_Q42 and Htt_ex1_Q21 root tips were intact, as in GUS (Fig. 2B). As observed in leaf cells, Htt_ex1_Q63 root cells in the root apical meristem (RAM) area often contained aggregates while Htt_ex1_Q42, Htt_ex1_Q21, and GUS had only a few or no aggregates (Fig. 2F). These results indicate that mHtt also induces aggregation in RAM cells and affects root growth and root hair outgrowth.

**Fig. 2.**
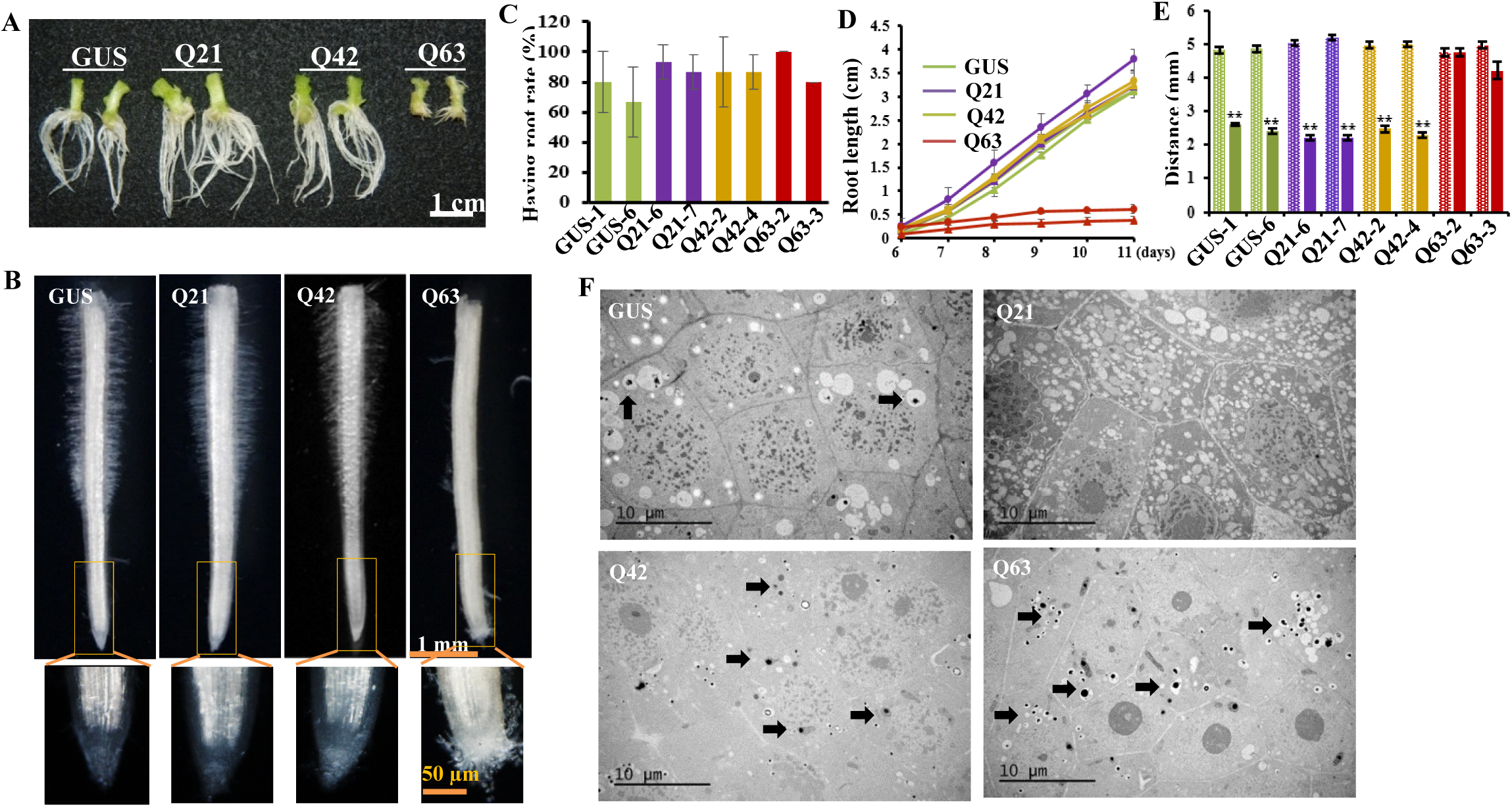
Morphology and microstructures of induced adventitious roots. (A) Root morphology. (B) Root hair distribution in the ∼5 mm root tip region and enlarged root tip structure. (**C, D)** Root emerging frequencies and root growth rates in subcultured shoots. Five shoots per line with two lines per genetic cassette were sub-cultured. The root emerging frequency was recorded at day 6 of subculture (C). The length of the longest root per plant was measured daily from day 6 till day 11 (D). The experiment was repeated three times. Both data plotted are the average of three independent experiments ± SD. **(E)** Root hair distribution. The full length of the longest induced root within 4-6 mm from each subcultured shoot and the distance between the root tip and root hair emerging site were measured. Data plotted are the average of all observed roots ± SD (GUS: n = 29 and 28; Htt_ex1_Q21s: n=25 and 25; Htt_ex1_Q42: n=26 and 28; Htt_ex1_Q63: n=21 and 29). **: *p* <0.01 level. **(F)** Root cells at the root apical meristem (RAM) area from four transgenic lines observed under TEM. Arrow indicates protein aggregates. Q21: Htt_ex1_Q21; Q42: Htt_ex1_Q42; and Q63: Htt_ex1_Q63.

### PolyQ63 remodels the proteome of roots

The significant differences in morphology and protein aggregation observed between Htt_ex1_Q63 and Htt_ex1_Q21 roots gave us an ideal pair of specimens for investigating toxic assault of polyQ- induced protein aggregation on cellular pathways. To gain insight into polyQ63-induced changes, a quantitative proteomic analysis was performed on Htt_ex1_Q63 and Htt_ex1_Q21 young roots (∼0.5 cm). The proteomic results (Text S1; Fig. S5; Tables S1-S3) are summarized in Fig. 3A. 2D hierarchical clustering analysis of 5,073 proteins quantified by ≥ 2 peptides (Table S4) indicates that extensive proteome remodeling occurred in Htt_ex1_Q63 roots (Fig. 3B). Using the cutoff of abundance change ≥1.5-fold and an FDR-corrected *p* <0.05, 1,693 proteins were identified as DAPs with 854 abundance-increased and 839 abundance-decreased (Table S5). Gene Ontology (GO) analysis showed that polyQ63 mainly suppressed proteins involved in translation, folding, transport and localization, especially from the vesicle-mediated ER to the Golgi (Text S1; Fig. S6; Table S6). It also induced a large group of proteins associated with oxidative stress (Fig. S7; Table S7). Based on molecular function, polyQ63 primarily perturbed GTPases, RNA binding proteins and enzymes involved in C_1_ group transfer (Fig. S6). In addition, 43 out of 49 mitochondria-related DAPs (Tables S8 and S9) were found to be abundance-increased. We, therefore, measured ATP levels in all four transgenic lines and detected significantly higher ATP levels in Htt_ex1_Q63 than the other three lines (Fig. S8). A large group of identified abundance- increased mitochondria-related DAPs along with higher ATP levels observed in Htt_ex1_Q63 indicates that cellular response to stress induced by polyQ63 is similar to what has been observed in young HD mice with elevated levels of glycolysis and ATP^27^, but different from those in old HD mice with dysfunctional mitochondria and low ATP^19, 28^. Our results revealed that polyQ63 extensively altered the root proteome and affected multiple cellular pathways as observed in other HD models^23, 29^.

**Fig. 3.**
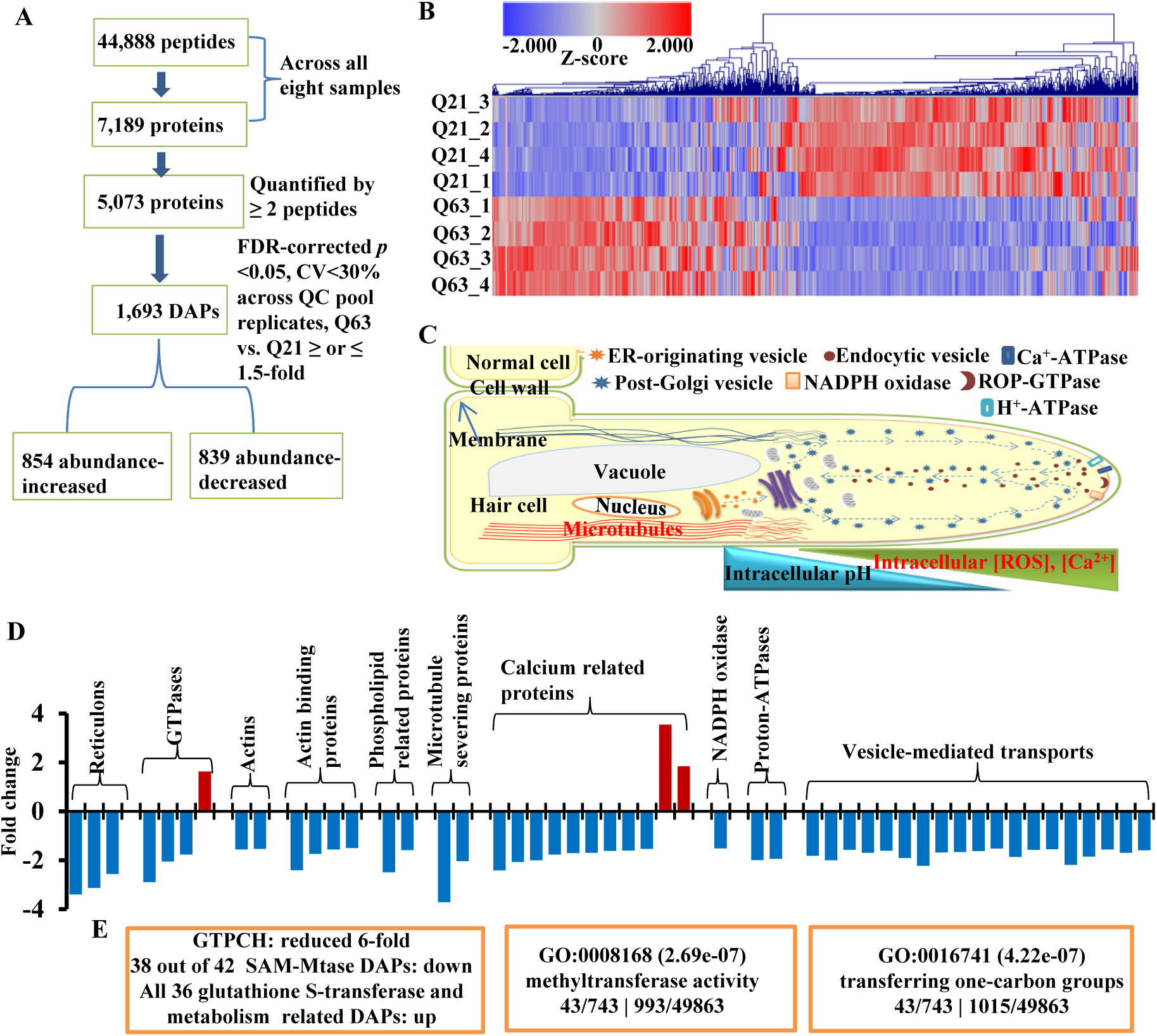
Proteome of induced adventitious roots. **(A)** Proteomic comparison between Htt_ex1_Q63 (Q63) and Htt_ex1_Q21 (Q21) roots (n = 4). **(B)** 2D hierarchical clustering analysis of the expression levels of 5,073 proteins quantified by ≥ 2 peptides. **(C)** A schematic representation of a root hair cell showing essential cellular components for its elongation adopted from Balcerowicz et al.^31^. **(D)** Obtained DAPs involved in root hair elongation. **(E)** Affected major proteins and pathways (orange boxes) related to C_1_ metabolism.

### PolyQ63 affects key proteins involved in the root hair tip-growth

Root hair tip growth resembles neuronal axon extension. There is continuous exocytosis of plasma membrane vesicles toward the tip, involving reconfiguration of the actin cytoskeleton and activation of actin-binding proteins and their upstream signals, such as calcium, phospholipids, and small GTPases (Fig. 3C)^30, 31^. In animals, GTPases interact with tubule-shaping proteins, reticulons and DP1/Yop1p, to form the tubular ER network for neurite elongation^24^. Our proteomics data showed that 47 out of 50 identified tubular ER network-related DAPs, including three reticulon-like proteins and three GTPases, were abundance-decreased (Fig. 3D; Table S10), supporting impaired root growth and root hair tip-growth observed in Htt_ex1_Q63. However, root hair initiation was unaffected (Text S2), as evidenced by a 1.5-fold decrease in GLABRA2 (GL2) (Fig. S9B), a key inhibitory transcription factor controlling the differentiation of an epidermal cell into a root hair cell^30, 31^. This is also supported by induced adventitious roots from Htt_ex1_Q63 shoots with root hairs when they were inoculated on the surface of the maintenance medium (Fig. S9C).

A root tip can be divided into meristematic, transition and elongation regions based on their specialized functions and characteristic cellular activities (Fig. S10A)^31^. Our proteomic data helped us to understand stunted root growth and deformed root apex. We identified a large group of abundance-decreased DAPs that function in cell division, expansion and elongation (Fig. S10B) and ribosome biogenesis along with a group of ribosome assembly factors, glutathione-*S*- transferases (GSTs), DNA-directed RNA polymerases, serine/threonine-protein kinases, and phosphatases (Fig. S10C; Tables S11-14). A decrease in their abundances may be responsible for defective root growth and root cap integrity, as discussed in Supplementary Text S3.

### PolyQ63 also affects proteins whose mammalian homologs play critical roles in NDDs

A group of DAPs affected by polyQ63 whose mammalian homologs are well-known players in NDDs were identified (Table 1). GTPCH was the most decreased enzyme (6-fold decrease) along with another member (UnitProt: W8TFR1, 2.5-fold decrease). In mammals, it plays an important role in the central nervous system by regulating the biosynthesis of BH_4_, which acts as a cofactor for the production of several monoamine neurotransmitters^17^. GTPCH has not been reported to be directly involved in HD before. In humans, GTPCH deficiency was found to deplete dopamine with an increased risk of Parkinson’s disease^32^. The next was a huntingtin-interacting protein K (HYPK)-like (2.5-fold decrease) whose mammalian homolog is an HD hallmark protein preventing mHtt-induced aggregation and apoptosis^33^. The third one was RHD3, a GTP-binding protein essential for root hair growth^34^, which decreased by 1.8-fold. It is a homolog of human atlastin-1^24–26^ whose mutation causes hereditary spastic paraplegia^24^. Moreover, a K Homology (KH) domain-containing protein whose human homolog is involved in schizophrenia^35^ was also reduced by 3.2-fold. In addition, 10 glutamate/GABA cycle-related enzymes^36^ were found to be increased (Table 1), suggesting that polyQ63 also alters the glutamate/GABA-glutamine cycle in transgenic plants.

**Table 1.** DAPs homologous to mammalian proteins associated with NDDs and involved in glutamate, GABA and glutamine metabolism.

| UniProtKB | Protein description | Log2 fold-change | p-value* |
| --- | --- | --- | --- |
| A0A1S3WYB9 | GTP cyclohydrolase 1 | -2.64 | 0.002 |
| W8TFR1 | GTP cyclohydrolase II | -1.32 | 2.5E-05 |
| A0A1S4DLE6 | Huntingtin-interacting protein K-like | -1.34 | 0.001 |
| A0A1S4DQ39 | Protein ROOT HAIR DEFECTIVE 3 homolog | -0.82 | 4.9E-05 |
| A0A1S4CML3 | KH domain-containing protein At1g09660/At1g09670-like | -1.70 | 6.3E-05 |
| **Glutamate, GABA and glutamine metabolism** |  |  |  |
| A0A1S4DRP0_TOBAC | Gamma aminobutyrate transaminase 3, chloroplastic-like isoform X1 | 1.79 | 1.3E-05 |
| A0A1S3ZU19_TOBAC | Glutamate dehydrogenase A-like | 1.08 | 5.7E-05 |
| A0A1S4AZM4_TOBAC | Glutamate--glyoxylate aminotransferase 2-like | 1.07 | 5.0E-05 |
| A0A1S4A4G8_TOBAC | Ferredoxin-dependent glutamate synthase 1, chloroplastic/mitochondrial-like | 0.89 | 4.5E-04 |
| A0A1S4AWH0_TOBAC | Glutamine synthetase | 0.84 | 5.1E-04 |
| Q7M242_TOBAC | Glutamate synthase (Ferredoxin) (Clone C(35)) (Fragment) | 0.83 | 6.6E-04 |
| A0A1S4AUA9_TOBAC | Bifunctional aspartate aminotransferase and glutamate/aspartate-prephenate aminotransferase-like | 0.83 | 1.0E-03 |
| A0A1S3Z1F3_TOBAC | Ferredoxin-dependent glutamate synthase, chloroplastic-like | 0.82 | 5.8E-04 |
| A0A1S4B8W9_TOBAC | Glutamate dehydrogenase | 0.82 | 3.1E-04 |
| A0A1S4AZA4_TOBAC | Gamma aminobutyrate transaminase 1, mitochondrial isoform X1 | 0.80 | 7.6E-04 |

\*: t-test with FDR correction

### PolyQ63 impairs C_1_ metabolic pathway enzymes

Importantly, our proteomic results also suggested alterations in C_1_ metabolic pathways in the Htt_ex1_Q63 transgenic roots (Fig. 3E), which comprises of the folate cycle, the methionine cycle, and the transsulfuration pathway^11, 12, 15^, with following evidences. First, GTPCH, a rate-limiting enzyme for plant folate biosynthesis (Fig. 4A)^37^ decreased 6-fold (Table 1), which could dramatically affect folate levels and impair C_1_ metabolism. Next, Htt_ex1_Q63 expression decreased the abundances of 38 out of 42 *S*-adenosyl methionine-dependent methyltransferases (SAM- MTases) (Table S15) and increased the abundances of all 36 GSTs and proteins involved in the regulation of glutathione redox status (Table S12). SAM-MTases play critical roles in the transfer of methyl groups to various biomolecules, while GSTs regulate redox status^11, 12^. These results imply that polyQ63 disturbs the methionine cycle and transsulfuration pathways of C_1_ metabolism. GO analysis of abundance-decreased DAPs also showed significant enrichment of methyltransferase activity and C_1_ transfer pathways (Fig. 3E; Fig. S6). In summary, our results showed that polyQ63 impairs GTPCH and many other proteins associated C_1_ metabolism in the young roots. These findings are novel and valuable for understanding mHtt-induced early pathophysiological changes since GTPCH is a rate-limiting enzyme for de novo folate biosynthesis in plants. Moreover, folate-mediated C_1_ metabolism is essential for cell survival and proliferation in both animals and plants by providing C_1_ units to various biosynthetic and methylation processes^11–16^.

**Fig. 4.**
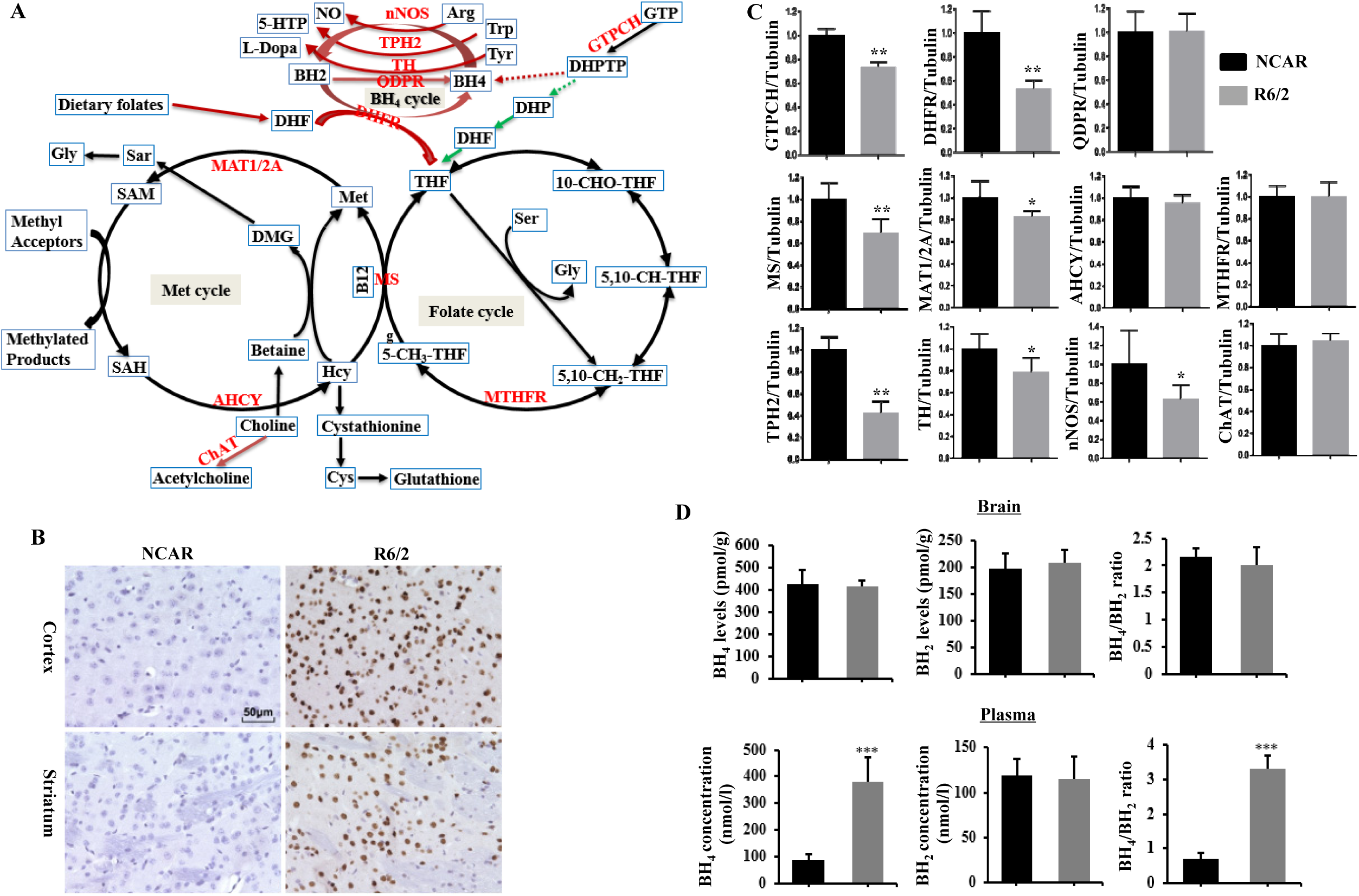
C_1_ metabolic pathway and characterization of R6/2 mice. **(A)** Folate, Met and BH_4_ cycles in plants and animals and their associated metabolism. Red lines stand for mammal specific, green lines stand for plant specific while black lines stand for both. All enzymes with protein levels examined by immunoblotting are marked in red. **(B)** mHtt protein aggregates in cortex and striatum regions were detected with anti-Htt antibody (mEM48) in four-week-old male R6/2 and NCAR mice. **(C)** Quantification analysis of immunoblotting results of GTPCH, DHFR, QDPR, MS, MAT1/2A, AHCY, MTHFR, TPH2, TH, nNOS and ChAT (n = 7). The band intensity of each protein from Western blotting was normalized with γ-tubulin on the same blot (Fig. S12). The ratio was further calculated against NCAR whose relative expression level was set as 1. All data plotted are the average (n = 7) ± SD. **(D)** Contents of BH_4_ and BH_2_, and their ratio in brain tissues and plasma (n = 4, average ± SD). *: *P*<0.05; **: *P*<0.01. ***: *P*<0.001. Abbreviations used for enzymes: AHCY, *S*-adenosylhomocystein hydrolase; ChAT, choline acetyltransferase; DHFR, dihydrofolate reductase; GTPCH, GTP cyclohydrolase I; MAT1/2A, methionine adenosyltransferase; MS, methionine synthase; MTHFR, methylene-tetrahydrofolate reductase; nNOS, neuronal nitric oxide synthase; QDPR, quinoid dihydropteridine reductase; TH, tyrosine hydroxylase (Tyr); TPH2, tryptophan hydroxylase.

### mHtt impairs GTPCH and DHFR in young HD mice

We then set out to validate whether the novel findings of mHtt-impaired GTPCH expression and C_1_ metabolic pathways in Htt_ex1_Q63 plants are true in R6/2 mice, a well-established HD animal model^38^. GTPCH catalyzes the same reaction involving the conversion of GTP to dihydroneopterin triphosphate (DHPTP) in both mammals and plants^37, 39^. However, the DHPTP produced in plants is converted into dihydrofolate to enter the folate cycle, while it is converted into BH_4_ in mammals to serve as a cofactor for several hydroxylases, which produce monoamine neurotransmitters with concomitant oxidation to dihydrobiopterin (BH_2_) (Fig. 4A). BH_2_ is then rapidly reduced back to BH_4_ by quinoid dihydropteridine reductase (QPDR) or DHFR when QPDR is limiting^40^. The main function of DHFR in animals is to generate a metabolically active form of tetrahydrofolate (THF) from diet-derived folic acid to enter the folate cycle^41^ while its reductive function also interconnects folate metabolism to biopterin metabolism^40^. Hence, we analyzed both GTPCH and DHFR, as well as nine C_1_ and BH_4_ metabolism-related enzymes, in the brains of 4-week-old male R6/2 mice (Fig. 4A).

In HD, the most affected organ is the brain, especially the striatum and the cortex^4, 21^. Four- week-old R6/2 mice (Fig. S11) showed no differences in body weight, brain weight or striatum size compared to the noncarrier (NCAR) control animals (Fig. S11B-S11E), but did display mHtt- containing protein aggregates in both the striatal and cortical regions (Fig. 4B). Notably, western blotting results showed that R6/2 brain tissues had significant decrease in the expression levels of GTPCH and DHFR, as well as of methionine synthase (MS) and methionine adenosyltransferase (MAT1/2A), but no change in the expression levels of QPDR, methylene-tetrahydrofolate reductase (MTHFR), or *S*-adenosylhomocysteine hydrolase (AHCY) (Fig. 4C; Fig. S12). In addition, the levels of all three BH_4_-dependent enzymes tryptophan hydroxylase (TPH2), tyrosine hydroxylase (TH) and neuronal nitric oxide synthase (nNOS) for the synthesis of serotonin, dopamine and nitric oxide were all significantly decreased, but not of BH_4_-independent choline acetyltransferase (ChAT) (Fig. 4C; Fig. S12). These results not only confirmed that the mHtt_ex1_- impaired GTPCH expression discovered in mHtt transgenic plant system is also true in young R6/2 mice, but also revealed mHtt_ex1_-impaired expression of DHFR at this juvenile stage.

### mHtt affects GTPCH and DHFR expression in striatum and cortex differently

Striatum and cortex were found to be affected by mHtt differently^4, 21, 42^ and transcriptional levels of *GCH1* (encoding for GTPCH) and *DHFR* were observed to be different in basal ganglia and cortex of the human brain^17^. Since GTPCH and DHFR are critical enzymes for BH_4_ biosynthesis and C_1_ metabolism, they were further analyzed in the striatum and cortex separately. Western blotting results showed that the expression of DHFR was significantly decreased in both regions while, surprisingly, the GTPCH level was decreased in the cortex, but increased in the striatum compared to those of NCAR control (Fig. 5A). The increase of GTPCH in the striatum and decrease in the cortex, and decrease of DHFR in both regions were further confirmed by immunohistochemistry assay (Fig. 5C-5F). These results indicated that GTPCH and DHFR are affected differently by mHtt_ex1_ in the striatum and cortex at an early stage.

**Fig. 5.**
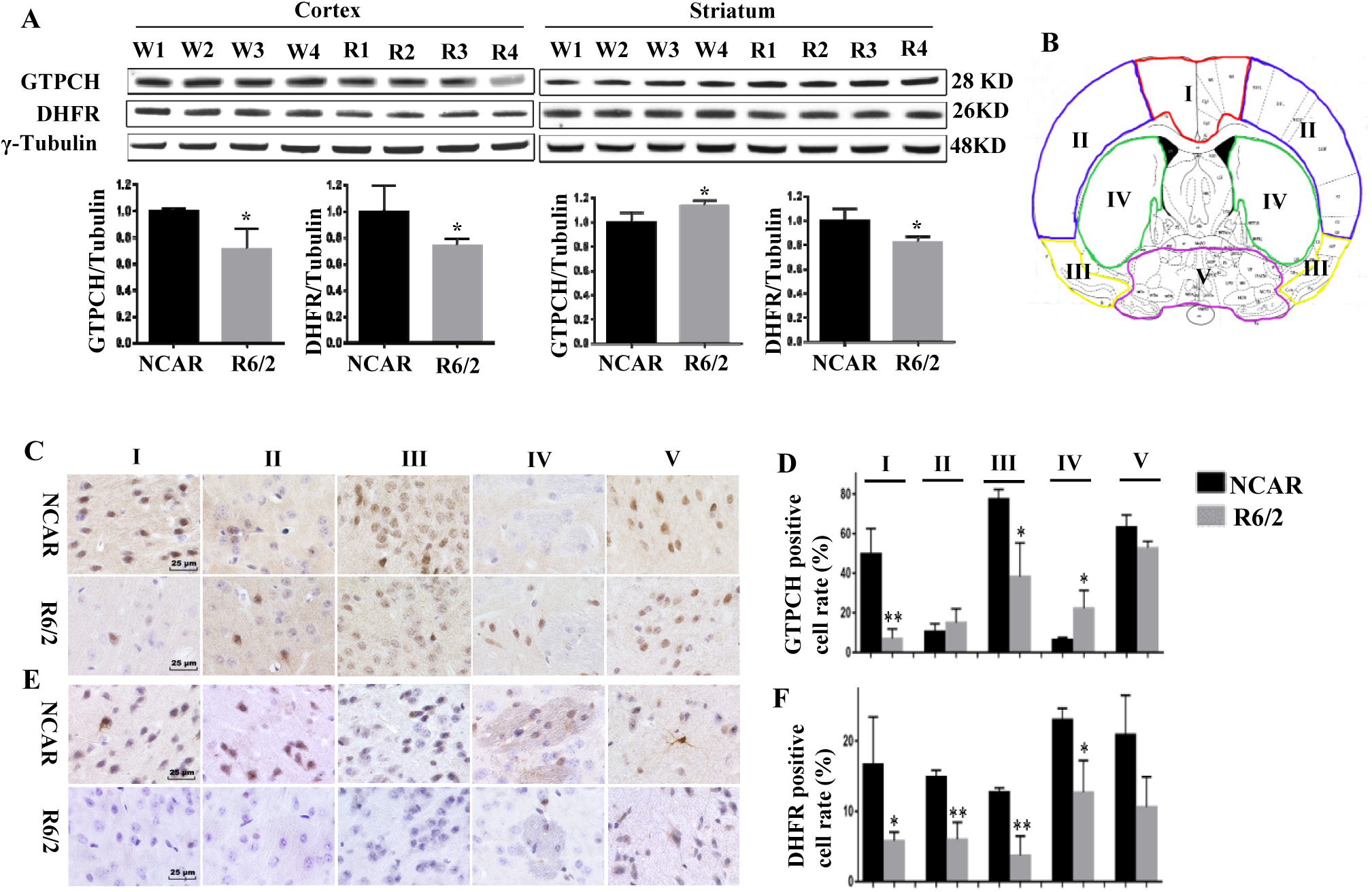
Western blotting and immunohistochemistry assay of cortex and striatum regions to quantify GTPCH and DHFR expressions. **(A)** Immunoblotting of GTPCH and DHFR in cerebral cortex and striatum tissues of R6/2 and NCAR mice using γ-tubulin as internal control (n=4). Protein bands were quantified as described in Fig. 3. **(B)** Diagram of mouse brain section divided into five regions: three regions of cortex (I-III); striatum (IV); hypothalamus and pallidum (V). **(C, D)** GPTCH and **(E, F)** DHFR positive cells and their ratios were quantified in five regions (I-V). *: *P*<0.05; **: *P*<0.01.

### mHtt alters C_1_ and BH_4_ metabolisms in young R6/2 mice

We further measured BH_4_, BH_2_ and free amino acid levels in both plasma and brain tissues of 4- week-old R6/2 mice to investigate any changes that may be related to impaired expression of GTPCH and DHFR. The results showed that BH_4_ was significantly increased by 4.5-fold in plasma but no change in brain tissues yet, while BH_2_ remained unchanged in both plasma and brain tissues (Fig. 4D). Although there was no reduction in BH_4_ in brain tissues, the dramatic increase in plasma could be the body’s response to its declining cerebral levels of BH_4_ since it can cross the blood-brain barrier^43^. The other reasons for no observed change in brain BH_4_ levels could be that residual enzymatic activity of GTPCH is still enough for BH_4_ biosynthesis or it might be compensated by the other alternate de novo or/and salvage pathways^17^.

We also analyzed free amino acids in both plasma and brain tissues with particular emphasis on Ser and Gly, as well as aromatic amino acids (Phe, Tyr, Trp). The former provide C_1_ units to various biomolecules while the latter serve as substrates for monoamine neurotransmitters. We detected 29 free amino acids in plasma and 26 in brain tissues (Table 2). We divided them into four groups based on their major involvement in the central nervous system: group I, sources of C_1_ units; group II, related to neurotransmitters; group III, branched-chain amino acids; and group IV, others. Among the five members in group I, Gly and Ser are the major sources of C_1_ units^44, 45^. Gly was increased significantly by 26.5% (*p* = 0.014) in the brain tissues, while Ser showed a marginally significant increase of 17.9% (*p* = 0.113) in plasma. In the folate cycle, Gly and Ser fuel mitochondrial enzymes through purine production^12, 15^. These results indicate that the C_1_ source in R6/2 mice was disrupted by mHtt_ex1_ at this early stage, which might potentially lead to mitochondrial dysfunction at the later stage.

**Table 2.**
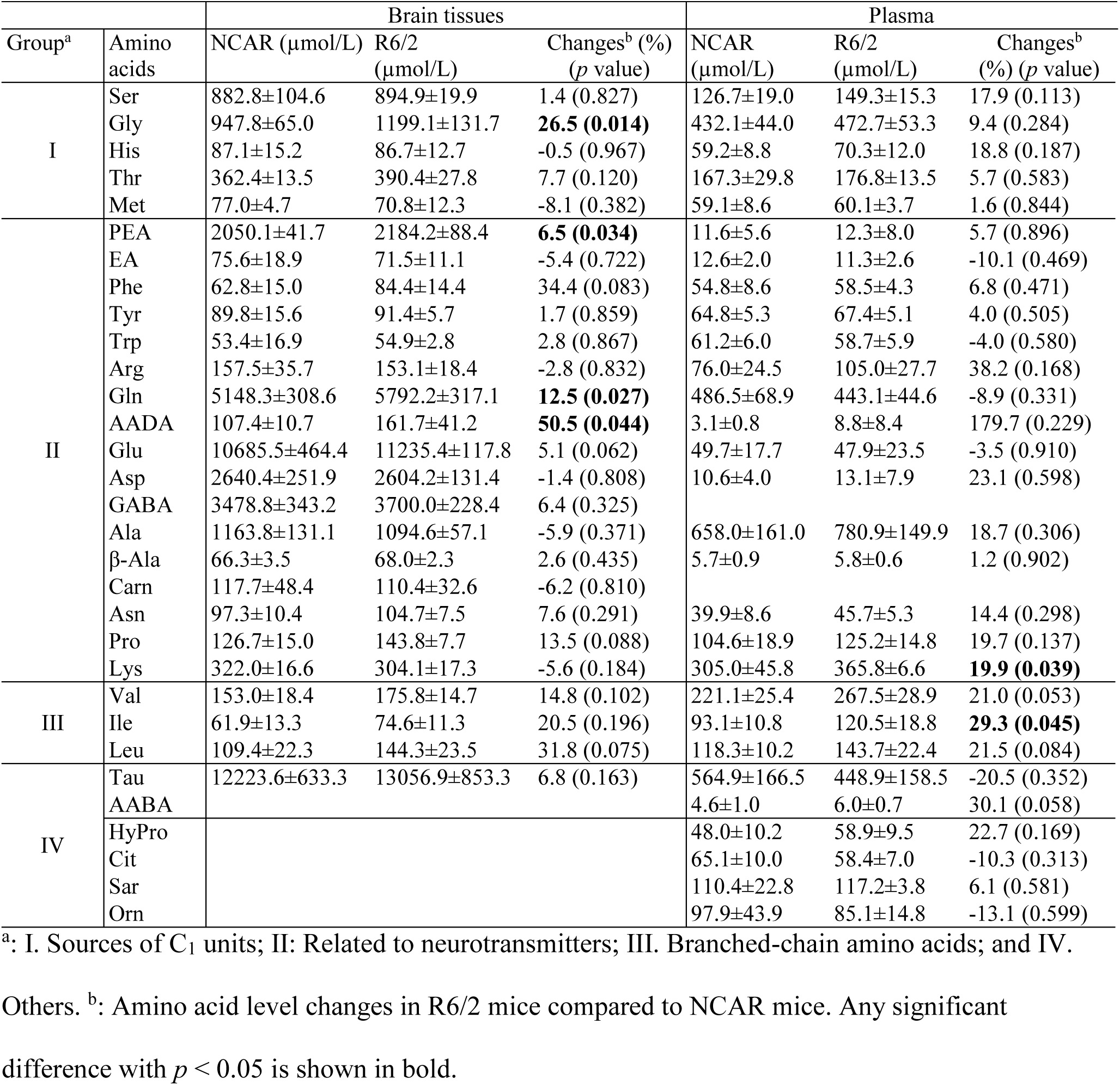
The contents of amino acids in R6/2 and NCAR mouse brain tissues and plasma.

Among group II amino acids involved in neurotransmission, PEA, Gln and AADA were found to be significantly higher (*p* <0.05), while Phe, Glu and Pro were less significantly higher (*p* <0.1) in brain tissues of R6/2 mice than NCAR control. In mammals, Phe is an essential amino acid and also serves as an important precursor of Tyr requiring BH_4_ as a cofactor for the synthesis of monoamine neurotransmitters^32, 39, 46^, while PEA regulates monoamine neurotransmission in neurons^47^. However, there was no change in the contents of three other monoamine neurotransmitter precursors Tyr, Trp and Arg in brain tissues yet. Concerning elevated Gln, Glu and Pro, they play important roles in the Gln/Glu (GABA) cycle^48, 49^. In regard to elevated AADA, it is a 6-carbon homologue of Glu and generated from the catabolism of Lys^50, 51^. Lys was significantly increased (∼20%) in R6/2 plasma. It is not clear whether there is any relation between high plasma level of Lys and elevated AADA content in brain tissues, the elevation of the latter in the brain tissues could exert its toxicity via enhanced release or transport of excitatory Glu^50, 51^. Undeniably, the increase of these amino acids along with the decreased levels of three BH_4_-dependent neurotransmitter synthases (Fig. 4C) implies that neurotransmitter biosynthesis or metabolism in 4-week-old R6/2 mice was perturbed by mHtt_ex1_.

Group III comprises of three proteinogenic branched-chain amino acids (Ile, Leu and Val), which are essential to build muscle^52^. They were reported to be reduced in the plasma of the middle- to aged-HD patients and sheep and thus proposed to be potential biomarkers of HD because their reductions were correlated with weight loss and disease progression^52–54^. In the current study, they were all substantially increased in both brain tissues and plasma of young R6/2 mice. The increase in these amino acids in the early stage could be a stress response to counteract the toxic effects of mHtt and was consistent with no body weight loss in young R6/2 mice (Fig. S11B). The contradiction between our results and previous reports implies that mHtt-induced metabolic dysfunction in the early growth stage is different from that in the middle and late growth stages. Nevertheless, elevated levels of amino acids and their derived products have been reported as pathogenic factors for neurological disorders^55^. The observed changes in plasma BH_4_ and the above amino acids indicate that mHtt_ex1_ alters BH_4_ and C_1_ metabolism in young R6/2 mice.

## Discussion

Although plants have not been considered previously to be used to study NDDs, our studies demonstrate that the plant-based system with polyQ-induced protein aggregation is unique and valuable for studying expanded polyQ-mediated toxicity. In the current study, our approaches involved establishing stably transgenic tobacco plants carrying *mHtt_ex1_*, observing polyQ length-dependent toxic effects, discovering mHtt-affected proteins and cellular pathways, and finally verifying novel findings from the plant-based system in young R6/2 mice. In transgenic plants, we observed that the toxic effects of expanded polyQ on protein aggregation and plant growth, especially root and root hair development, resemble those reported in animal HD models in a polyQ length-dependent manner^18, 19^. Importantly, using this plant-based system allowed not only the identification of many homologous proteins known to be affected by expanded polyQ in neuronal cells but also the discovery of hitherto unreported mHtt-mediated disturbance of GTPCH. Impaired GTPCH expression and its associated C_1_ metabolism could be an early event in the pathogenesis of mHtt. These findings recognize autotrophic plants with de novo synthesis of folate^11, 12^ to be a good system for studying mHtt-impaired folate and C_1_ metabolic pathways.

Because of the evolutionary distance between plants and mammals, it is essential to validate mHtt-impaired GTPCH expression and C_1_ metabolism in an HD animal model to confirm the usefulness of the plant-based polyQ system. In animals, GTPCH controls the biosynthesis of BH_4_^39^, an essential cofactor for amino acid hydroxylases to produce monoamine neurotransmitters^32, 39^ while DHFR plays dual roles in the folate and BH_4_ cycles (Fig. 4A)^40, 41^. Our validation studies confirmed that the expression levels of both GTPCH and DHFR were suppressed in total brain tissues (Fig. 4C). Unexpectedly, using plant and animal HD systems together led to the discovery of impaired expression of GTPCH and DHFR, which have not been directly linked to HD before. Only recent genome-wide association studies with genetic information from a large number of HD patients predicted that DHFR is possibly a modifier of disease progression^5, 56^. In addition, genetic defects in DHFR were found to cause insufficient cerebral BH_4_ levels^40^, while GTPCH-deficient mice exhibited BH_4_-related metabolic disturbance and infancy-onset motor impairments^57^. These previous reports point to the importance of maintaining normal expression of GTPCH and DHFR in animals. The current study provides direct evidence that mHtt impairs these two key enzymes of C_1_ and BH_4_ metabolism. The toxic effects of mHtt on GTPCH and DHFR are also supported by the observed interruption of C_1_ and BH_4_ metabolism in young R6/2 mice. Although there was no decrease in BH_4_ levels in brain tissues yet, the dramatic increase in plasma BH_4_ levels and the decrease in three BH_4_-dependent aromatic amino acid hydroxylases imply declining cerebral levels at this young stage (Fig. 4D). In addition, we observed elevated levels of several amino acids serving as major sources of C_1_ units or precursors for neurotransmitter synthesis (Table 2). Taken together, our study revealed that GTPCH- and DHFR-associated C_1_ and BH_4_ metabolism and neurotransmitter biosynthesis were possibly disturbed by mHtt at a very early stage.

In the case of inherited polyQ diseases, by the time any disease symptoms appear, polyQ- induced cellular damage might already be too advanced for treatment to be successful. Any strategy preventing the development of diseases or impeding the progression of diseases could be a promising avenue for improving health and quality of life for mutant gene carriers. Our findings of early ployQ-affected GPTCH and DHFR, and their related cellular pathways are significant because disordered transcriptomes and altered epigenetics have been reported to be common HD- induced pathological changes^6, 58, 59^. Decreased DHFR levels by mHtt observed in the current study provide a clue to understanding how mHtt affects C_1_ metabolism and the flux of methyl groups, leading to potentially low methylation activity, and subsequently causing transcriptional dysregulation and epigenetic alteration. In plants and mammals, the folate metabolic pathway generates and provides methyl groups for methionine biosynthesis from homocysteine. More than 80% of synthesized methionine is further converted to SAM, which is a universal methyl donor for methylation reactions leading to the synthesis of many biomolecules and epigenetic regulation of gene expression via DNA and histone methylation^11, 12^. In addition, the observed opposite expression of GTPCH in the cortex and striatum in young R6/2 mice (Fig. 5) are also notable.

Under healthy conditions, the cortico-striatal system is tightly regulated by dopamine^42, 60^. mHtt- mediated opposite expression of GTPCH in the striatum and cortex may cause imbalance of the cortico-striatal system through dysregulation of dopamine and contribute to the selective vulnerability of striatal medium-spiny neurons to toxic polyQ repeats^42, 60^. Aberrant communication between striatum and cortex has been considered to play a critical role in striatal neuronal loss^4, 42^. It is not clear at this time how mHtt_ex1_-mediated protein aggregation affects GTPCH expression in plants and animals. Created Htt_ex1_Q63 transgenic plants with dramatic changes in the abundances of GTPCH (Table 1) are valuable. They together with HD animal models could be used to further understand the relationship between the neurotoxicity and early polyQ-mediated the alterations of C_1_ and BH_4_ metabolisms, epigenetics and transcriptomes to identify therapeutic targets and develop preventive strategies for the HD.

In addition, neuronal loss is a common pathological hallmark of HD and many other NDDs^1, 2^. Root hairs and neuronal cells share some strikingly similar characteristics, such as polarity, long extension outgrowth, high energy demand, and environmental sensing capacity^7, 8^. Observation of restricted root hair out-growth in Htt_ex1_Q63 roots and partially restricted in Htt_ex1_Q42 roots means that root hairs might also respond to toxic polyQ repeats similar to neuronal cells. In particular, many affected components of tubular ER network that were identified in Htt_ex1_Q63 roots (Fig. 3D; Table S10) are known to play crucial roles in neurite elongation. These results suggest that toxic polyQ repeat-induced neuronal loss and restricted root hair out-growth may share some common mechanism(s). With recent seminal discovery in adult neurogenesis that the adult brain may be able to make new neurons throughout adulthood^61^, our results that expanded polyQ repeat restricted root hair out-growth (Fig. 2B and 2E), but did not affect hair initiation (Fig. S9B and S9C) may provide another piece of useful information for future polyQ-mediated NDD study. While many study the mechanisms of expanded polyQ-induced age-dependent progressive neurodegeneration, we should pay attention to the toxic effects of polyQ repeats on neuronal development and neurite outgrowth.

In summary, our studies demonstrated for the first time, to the best of our knowledge, that expanded polyQ63-triggered protein aggregation in plant cells is akin to that in mammalian cells with many affected plant proteins homologous to mammalian ones, which play critical roles in NDDs. The obtained results from both plant and animal studies proved the concept that the plant-based protein aggregation system is useful for studying abnormal polyQ-mediated cellular toxicity in general and dysregulation of C_1_ metabolism in particular. Given the importance of C_1_ metabolism and the BH_4_ cycle^13, 15, 39^, the observed changes in GTPCH, DHFR, and certain C_1_/BH_4_ metabolism-related enzymes and metabolites in young HD mice could be major contributing factors to HD onset and progression. Our findings from plant-based and R6/2 HD models open a new avenue to study the roles of GTPCH, DHFR, and C_1_ and BH_4_ metabolism in the initiation and progression of HD, and perhaps of other polyQ diseases.

## Materials and methods

### Plant materials and transgenic plant maintenance

Tobacco (cultivar “W38”) plants were used to generate transgenic plants. Leaf explants from 4- week-old young plants prepared from seedlings were used for transformation. Methods of seed sterilization and leaf explant transformation were the same as described previously^62^. All PCR-confirmed transgenic plants were propagated and maintained on a maintenance medium (MS medium supplemented with 100 mg/L Timentin) under reduced light conditions of ∼15 µmol m^-2^ s^-1^ photoperiod (16 h d^-1^) at 23°C because Htt_ex1_Q63 plantlets showed stress symptoms with yellowish colored leaves and slow growth when grown under ∼60 μmol m^-2^ s^-1^ light intensity. For each genetic cassette, seven independent transgenic lines were selected for propagating genetically identical plants from each line for downstream study. To achieve that, a shoot (∼3 cm) was first cut and placed in MS medium for rooting. The remaining base part of plant with several nodes was continually subcultured for two weeks to obtain additional new shoots. Propagated plants from each transgenic line were used for downstream studies. Transgenic plants grown in soil were also maintained under reduced light intensity conditions.

### Construction of genetic cassettes

Three DNA sequences having 283, 346 and 409 bp encoding truncated Htt *N*-terminal fragments with 21Q, 42Q and 63Q repeats (GenBank Accession #: MK291497-MK291499) were synthesized (Life Technologies) by adding *Xba*I and *Sac*I restriction enzyme cutting sites at the 5’ and 3’ ends of each DNA sequence for sub-cloning. They were cloned into the plant expression vector pBI121 by replacing the bacterial *uidA* gene. Resultant three genetic cassettes were named as Htt_ex1_Q21, Htt_ex1_Q42 and Htt_ex1_Q63, which were introduced into *Agrobacterium tumefaciens* strain LBA4404 using the freeze-thaw method^63^.

### *Agrobacterium*-mediated transformation

Among plants, the tobacco plant is well suited for this study because of its well-studied physiology and biochemistry, ease of genetic engineering with high transformation efficiency, and sufficient duplication of identical transgenic plants for various downstream studies^64, 65^.

*Agrobacterium*-mediated leaf disc transformation^66^ was employed for creating transgenic plants with the same protocols for *Agrobacterium* infection, culture media for induction and selection of kanamycin-resistant shoots, and rooting of kanamycin-resistant plantlets as described previously^62^. To study the transformation efficiencies of three genetic cassettes, Htt_ex1_Q21, Htt_ex1_Q42 and Htt_ex1_Q63, approximately 20 leaf discs were used for each cassette. The original pBI121 plasmid DNA with *uidA* gene encoding β-glucuronidase (GUS, designated as “GUS”) was used as a transformation control. For statistical analysis, 20 infected leaf discs were evenly cultured onto four selection plates, and each plate was calculated as one replicate. Infected leaf discs were transferred onto new media every two weeks. At the end of the second transfer, the numbers of induced calluses and shoots were recorded. The total transformed response was calculated as the number of responding explants (either with kanamycin-resistant calluses, shoots, or both) divided by the total inoculated explants. The transformation efficiency experiment was repeated three times. All leaf discs were maintained in a growth chamber under ∼60 μmol m^-2^ s^-1^ light intensity at 25°C.

### Genomic DNA PCR analysis

To confirm the presence of integrated *Htt_ex1_* with different CAG repeats and the kanamycin resistance gene *nptII* in kanamycin-resistant plants, PCR amplifications were performed. To detect *Htt_ex1_* with different CAG repeats, a pair of primers HDJamaF: 5’- ATGAAGGCCTTCGAGTCCCTCAAGTCC-3’ and HDJamaR2: 5’-CGGCGGCGGCGGTGGCGGCTGTT-3’^67^ was used. For detecting *nptII*, a pair of primers, NPT-II 5’: 5’-GTGGATCCCGCATGATTGAA-3’ and NPT-II 3’: 5’-TCGGATCCCTCAGAAGAACT-3’, was used. For each construct, seven independent transgenic lines were analyzed. Total genomic DNA was isolated from young leaves using a DNeasy® Plant min Kit (Qiagen). The DNA concentration was measured using an ND-1000 spectrophotometer (NanoDrop Technologies). For *nptII* detection, each 25 µl reaction mixture contained 100 ng DNA template, 2.5 µl of 10x PCR buffer, 1.25 units of Taq DNA polymerase (Sigma-Aldrich), and final concentrations of 0.2 mM dNTPs, 300 nM of each primer and 2 mM MgCl_2_. PCR cycles consisted of an initial denaturing step of 94°C for 2 min, followed by 30 amplification cycles of denaturation at 94°C for 1 min, annealing at 55°C for 60 s and elongation at 72°C for 30 s, and a final extended elongation at 72°C for 10 min. To amplify *Htt_ex1_* with different CAG repeats, each 30 µl reaction mixture contained 100 ng DNA template, 3 µl of 10x homemade PCR buffer^68^, 1.5 units of Taq DNA polymerase (Sigma-Aldrich), and final concentrations of 0.25 mM dNTPs, 300 nM of each primer and 2 mM MgCl_2_. PCR cycling conditions consisted of an initial denaturing step of 96°C for 3 min, followed by 35 amplification cycles of denaturation at 96°C for 45 s, annealing at 64°C for 45 s and elongation at 72°C for 1 min, plus the final extended elongation at 72°C for 10 min. PCR was performed on a TGradient Thermocycler (Biometra). Amplified DNA was resolved on a 1.0% agarose gel and visualized under UV light using UVP GelDoc-It™ Imaging Systems (Analytik Jena).

### RT-PCR analysis

To detect *Htt_ex1_* transcripts with different CAG repeats in transgenic plants, RT-PCR was used. Total RNA was isolated from young leaves using an RNeasy Plant Mini Kit (Qiagen). First strand cDNAs were made using a High-Capacity cDNA Reverse Transcription kit (Applied Biosystems) according to the manufacturer’s instructions. For RT-PCR, all PCR conditions were the same as for genomic DNA amplification except cDNA amplified from 100 ng of RNA was used instead of DNA template.

### Observation of root growth and morphology

For monitoring root growth, two independent lines from each genetic cassette were used. For each line, five propagated identical young shoots ∼3 cm in length were subcultured in the maintenance medium under a light intensity of ∼15 µmol m^-2^ s^-1^ with a 16 h d^-1^ photoperiod at 23°C. After subculture for six days, the number of plants showing emergence of roots was recorded.

Meanwhile, the length of its longest root among all emerged roots from each plant was measured every 24 h from day 6 to day 11. The experiment was repeated three times.

To observe the roots and root tips, a stereomicroscope with a Digital Sight DS-FI1 camera (Nikon) and NIS Elements version 3.2 software were used. About 21 to 29 propagated shoots per transgenic line were used and subcultured under the above conditions. After 7 to 9 days of subculture, when roots reached ∼5 mm in length, only the longest root per subcultured shoot was excised and gently cleaned to remove agar for photographing. First, it was placed on a microscopic slide with a few drops of water to create adhesion. Then it was covered with a cover slip to observe root length and root hair distribution and a picture was taken with a stereomicroscope at 1x magnification. The measuring tool in Adobe Photoshop software (Adobe Systems) was used to measure the total root length and the length from the root hair emerging site to the root tip.

### Electrophoresis and western blotting of protein extracts made from transgenic plants

Young leaves of transgenic plants grown in culture containers containing maintenance medium were harvested and ground into fine powder in liquid nitrogen. For SDS-PAGE, 150 mg each of fine powder were mixed with 350 µl of extraction buffer [50 mM Tris-HCl pH 8, 150 mM NaCl, 1% (v/v) NP40, 0.5% (w/v) sodium deoxycholate monohydrate, 0.1% (w/v) SDS, 1 mM 2- mercaptoethanol, 1 mM PMSF, 0.5 mM DTT, 1% (v/v) Sigma-Aldrich plant protease inhibitor cocktail]. After vortexing, the mixture was kept on ice for 10 min, and then subjected to two rounds of 20,000 x g centrifugation for 10 min at 4°C. The collected supernatant was mixed with Laemmli sample buffer and denatured at 95°C for 5 min. The protein concentration was determined by the Bradford method. Approximately 15 µg of protein were separated on a 13.5% SDS-PAGE gel in running buffer containing 192 mM glycine, 25 mM Tris and 0.1% (w/v) SDS at 100 volts for 70 min until the 14 kD protein marker in SeeBlue Plus2 pre-stained protein standard (Life Technologies, USA) reached the bottom end of the gel. Following separation, proteins were transferred to PVDF membrane using NuPAGE (Invitrogen) transfer buffer containing 10% (v/v) methanol and NuPAGE antioxidant. To show equal protein loading, the membrane was stained with 0.2% (w/v) Amido Black in 10% (v/v) acetic acid and destained with H_2_O.

For BN-PAGE (Invitrogen) immunoblotting, 100 mg each of fine leaf powder was mixed with 330 µl of extraction buffer containing 18 mM 3-[(3-cholamidopropyl) dimethylammonio]-1- propanesulfonic acid (CHAPS) in TBS with 10 µg/ml DNase I, 2 mM MgCl_2_, and 1% (v/v) plant protease inhibitor cocktail (Sigma-Aldrich). After a short vortex, the mixture was incubated at 21°C for 30 min with intermittent mixing. It was then subjected to one round of 20,000 x g centrifugation for 20 min at 4°C and a second round for 5 min. The collected supernatant was mixed with Native PAGE sample buffer containing 1% (w/v) Digitonin and 0.25% (w/v) Coomassie G-250. The proteins were separated on a NativePAGE (Invitrogen) 3-12% Bis-Tris gel in a NativePAGE running buffer (only cathode buffer containing 0.002% Coomassie G-250) at 100 volts running into 1/3 of gel then 40 volts till dye front reaching the bottom end of the gel. Following separation, proteins were transferred to PVDF membrane using NuPAGE transfer buffer at 25 volts for 1 h. To visualize the unstained protein standard NativeMark (Invitrogen), the membrane was stained with 0.1% (w/v) Coomassie Brilliant Blue R-250 in 50% (v/v) methanol and destained with a solution containing 7% (v/v) acetic acid and 40% (v/v) methanol.

To detect target proteins, the membrane was blocked with 5% (w/v) BSA in PBST overnight at 4°C and probed with anti-Huntingtin (1:1,000, ab109115, Abcam), anti-Ubiquitin (1:200, U5379, Sigma-Aldrich, USA) and biotinylated anti-Hsp70 (1:10,000, ab183437, Abcam) in blocking buffer for 1 h at 25°C. The secondary antibody used for detecting anti-Huntingtin and anti-Ubiquitin was 1:10,000 anti-rabbit IgG conjugated with HRP. To detect biotinylated anti- Hsp70, 1:20,000 HRP conjugated Streptavidin (1 mg/ml) was used. Luminescent signals were generated after incubation with SuperSignal® West Pico Chemiluminescent substrate (Pierce Biotechnology) and captured with Kodak Biomax X-ray film (PerkinElmer).

### Filter retardation assay

Protein extracts used for BN-PAGE were diluted with 0.1% (w/v) SDS in PBS at 1:3 ratios. Samples containing 10 µg of proteins were filtered through Whatman Cellulose Acetate Membrane Filters with 0.2 µm pore sizes (GE Healthcare), which have very low protein binding capacity. Filtration was performed using an Easy-Titer® ELIFA System 77000 (Pierce Biotechnology) according to the manufacturer’s instructions. After washing twice with 0.1% (w/v) SDS in PBS, the membranes were blocked with 10% (w/v) skim milk in PBST for 2 h at 25°C. Then, the membranes were incubated with primary antibodies: 1:2,000 anti-Ubiquitin and 1:20,000 anti-Huntingtin overnight at 4°C. After washing three times with PBST, the membranes were incubated with 1:20,000 anti-rabbit IgG conjugated with HRP for 1 h at 25°C. To detect Hsp70, the membrane was blocked with 5% (w/v) BSA in PBST and then incubated with 1:20,000 biotinylated anti-Hsp70 overnight at 4°C, followed by 1:40,000 HRP conjugated Streptavidin as a secondary antibody. The luminescent signals were detected as described above. To visualize the proteins, membranes were stained with 0.1% (w/v) Coomassie Brilliant Blue R- 250 in 50% (v/v) methanol for 15 min and destained with a solution containing 7% (v/v) acetic acid and 40% (v/v) methanol.

### Transmission Electron Microscopy (TEM) analysis

To observe cell structure and any aggregates in transgenic plants, TEM analysis was performed at the North Carolina State University Center for Electron Microscopy. Newly developed young leaves and roots of Htt_ex1_Q21, Htt_ex1_Q42, Htt_ex1_Q63 and GUS transgenic plants grown in culture containers were used. Samples were cut into 1 mm^3^ blocks for leaves and 0.5 mm (length) for root tips and then fixed in 3% glutaraldehyde in 0.05 M KPO_4_ buffer (pH 7.0) at 4°C. Samples were post-fixed in 2% OsO_4_ in the same buffer at 4°C in the dark. After dehydration with a graded series of ethanol, they were infiltrated and embedded with Spurr’s resin (Ladd Research Industries). Samples were sectioned with a Leica UC6rt ultramicrotome (Leica Microsystems) and placed onto 200-mesh grids. The grids were then stained with 4% aqueous uranyl acetate in the dark at 25°C followed by three distilled water washes at 40°C and 1 min in Reynold’s lead citrate followed by three more distilled water washes. All sections were observed under a JEOL JEM 1200EX transmission electron microscope (JEOL USA Inc). Images were captured using a Gatan Erlangshen Model 785 ES1000W camera and Digital Micrograph software (Gatan Inc). On average, each grid had approximately ten squares containing approximately five observable cells.

### Quantitative proteomic analysis

#### Sample preparation for mass spectrometry analysis

Quantitative proteomic analysis was performed at the Duke Proteomics and Metabolomics Shared Resource. To prepare samples, propagated shoots from Htt_ex1_Q21-6 and Htt_ex1_Q63-3 lines ∼3 cm in length were subcultured to induce roots. After 7 to 8 days of subculture, induced adventitious roots shorter than ∼0.5 cm were harvested. Due to their small sizes, each sample of pooled roots with a weight of ∼100 mg was frozen together at -80°C as one biological sample. Four biological samples from each line were prepared, and protein extraction was performed as follows. Harvested roots were first ground in liquid nitrogen and then resuspended in 200 µl of a buffer containing 4% (w/v) SDS and 50 mM triethylammonium bicarbonate (TEAB), pH 8.5, followed by probe sonication and heating at 80°C for 5 min. An additional sonication step was applied. After that, the samples were centrifuged, and a BCA assay was performed on the supernatants to determine the protein concentration. Twenty-five micrograms of each sample was digested with trypsin at 1:25 (w/w) trypsin:protein using S-trap™ processing technology (Protifi). After lyophilization, peptides were reconstituted in 50 µl of 1% (v/v) trifluoroacetic acid/2% (v/v) acetonitrile (MeCN). For QC analysis, a QC pool was made by mixing equal quantities of all samples.

#### Quantitative mass spectrometry analysis

For quantitative mass spectrometry analysis, quantitative one-dimensional liquid chromatography coupled with tandem mass spectrometry (1D-LC-MS/MS) was performed with 2 µl of the peptide digests per sample in singlicate, including additional analyses of conditioning runs and three QC pools. The QC pooled sample was analyzed three times at the beginning and interspersed throughout the analysis of individual samples, which were alternated between the two treatment groups (Table S1). Samples were analyzed using a nanoACQUITY UPLC system (Waters) coupled to a Q-Exactive HF-X high resolution accurate mass tandem mass spectrometer (Thermo Fisher Scientific) with a nanoelectrospray ionization source. Briefly, the sample was first trapped on a Symmetry C18 column (180 µm × 20 mm) (5 μl/min at 99.9/0.1 v/v H_2_O/MeCN), followed by analytical separation using a 1.7 µm AQCUITY HSS T3 C18 column (75 µm × 250 mm) (Waters) with a 90 min gradient of 5 to 30% (v/v) MeCN containing 0.1% (v/v) formic acid at a flow rate of 400 nl/min and column temperature of 55°C. Data collection on the HF-X MS was carried out in data-dependent acquisition (DDA) mode with a 120,000 resolution (@ m/z 200) full MS scan from m/z 375 to 1,600 with a target AGC value of 3e6 ions and 50 ms maximum injection time (IT).

The top 30 peptides were selected for MS/MS using a resolution of 15,000, max AGC of 5e4 ions and minimum AGC target of 2.25e3, max IT of 45 ms, collision energy of 27, an isolation width of 1.2 m/z and 20 s dynamic exclusion. The total analysis cycle time for each sample injection was approximately 2 h.

#### Protein identification and quantitation

Following UPLC-MS/MS analyses, data were imported into Rosetta Elucidator v4.0 (Rosetta Biosoftware Inc), and analyses were aligned based on the accurate mass and retention time of detected ions (“features”) using the PeakTeller algorithm.

Relative peptide abundance was computed based on the area-under-the-curve of the selected ion chromatograms of the aligned features across all runs. The MS/MS data were searched against a custom Swiss-Prot/TrEMBL database with *N. tabacum* taxonomy (downloaded on 04/30/2018). To minimize sequence redundancy, the database was further curated using cd-hit (http://weizhongli-lab.org/cdhit_suite/cgi-bin/index.cgi?cmd=cd-hita) with a sequence identity cut-off of 0.9, and an equal number of reverse entries was added for decoy database searching. The final database had 90,193 total entries. Mascot Distiller and Mascot Server (v 2.5, Matrix Sciences) were employed to produce fragment ion spectra and to perform the database searches. The parameters used for the database search included a precursor mass tolerance of 5 ppm, a production mass tolerance of 0.02 Da, trypsin specificity with up to 2 missed cleavages, fixed modification on Cys (carbamidomethyl) and variable modification of *N*-terminal protein acetylation. Individual peptide was scored using the PeptideProphet algorithm in Rosetta Elucidator, and data were annotated at a 1.0% peptide false discovery rate. For quantitative analysis, the data were first curated to contain only high-quality peptides with appropriate chromatographic peak shapes and the dataset was intensity scaled to the robust median across all samples. To analyze abundance, the expression values of these 5,073 proteins quantified by 2 or more peptides were Z-score-normalized followed by 2D hierarchical clustering analysis in Rosetta Elucidator.

Meanwhile, both analytical variability and biological variability were analyzed to determine the quality of quantitative proteomic analysis. To assess technical reproducibility, the percentage (%) of coefficient of variation (CV) was calculated for each protein across the three injections of a QC pool that were interspersed throughout the study, while %CVs were also measured for each protein across the individual analyses to assess biological variability.

#### GO enrichment and heatmap visualization analyses

DAPs (1,693 total with 854 abundance- increased and 839 abundance-decreased) with abundance changes more than 1.5-fold were subjected to GO enrichment analysis using the AgriGO platform v2.0 webserver (http://systemsbiology.cau.edu.cn/agriGOv2/)^69^. The UniProt batch retrieval tool (http://www.uniprot.org/uploadlists) was used to map proteins to tobacco (*N. tabacum*, cv. TN90) proteome (ID: UP000084051) to obtain UniProtKB for GO term identification. Of the 73,605 proteins of the tobacco proteome, 49,741 had GO annotation. Among 1,693 DAPs, 1,366 had GO annotation. To expand the GO annotation of the remaining 327 DAPs, these protein sequences were further used to query the uniprot_proteome (ID: AUP000004994) of *Solanum lycopersicum* (Tomato) (Strain: cv. Heinz 1706) using the BLASTP program with an E-value ≤ 1e-5 and identity ≥ 40% as the cutoff. The results added GO annotation for additional 122 proteins, whereas 205 proteins remained unannotated. Overall, the GO annotation of 745 (698 from tobacco, 47 from tomato) abundance-increased proteins and 743 (668 from tobacco, 75 from tomato) abundance-decreased proteins were included in the GO enrichment analysis. The GO terms of the abundance-increased or abundance-decreased proteins were imported separately into AgriGO for GO enrichment analysis. AgriGO’s Singular Enrichment Analysis (SEA) was used to identify enriched GO with default settings [Statistical test method: Fisher, Multi-test adjustment method: Yekutieli (FDR under dependency), Significance Level: 0.05, Minimum number of mapping entries: 5, and Gene ontology type: complete GO]. Selected enriched GO terms with biological importance were presented by GO Enrichment Plot (EHBIO gene technology).

To show the increased or decreased levels of DAPs associated with root growth and the maintenance of root apex structure, a heatmap visualization analysis was performed using Heml version 1.0.1 software^70^. The value assigned to each protein in Htt_ex1_Q21 and Htt_ex1_Q63 was calculated first as the expression ratio by comparing the relative abundance of any individual protein with its average of the two treatment groups, and then subjected to log2 transformation as described^71^.

### Animals

All procedures for using animals in the experiments were carried out in accordance with the National Institutes of Health ‘Guide for Care and Use of Laboratory Animals’ and were approved by the Institutional Animal Care and Use Committee of North Carolina Central University. Three- week-old R6/2 (B6CBA-Tg(HDexon1)62Gpb/125J) HD mice expressing *Htt_ex1_* carrying approximately 120 +/- 5 (CAG) repeats and 3-week-old wild type B6CBA-Nocarrier mice (NCAR) were purchased from the Jackson Laboratory (Bar Harbor). The animal housing room had a controlled temperature of 20 ± 2°C, humidity of 40-60%, and a 12-hour light/12-hour dark cycle. Animals had free access to food (Rodent Diet 20, PicoLab) and water. Both types of mice at 4 weeks of age were used first to collect blood for metabolite analysis and then sacrificed to harvest the brain for western blotting, histopathology and immunohistochemistry.

### Electrophoresis and western blotting of protein extracts from brain tissues

To isolate proteins from total brain tissues (n = 7), whole brain frozen in liquid nitrogen and stored at -80°C freezer was used. The tissues were individually homogenized with N-PER™ Neuronal Protein Extraction Reagent (Cat. #: 87792, Thermo Fisher Scientific) containing Halt Protease and Phosphatase Inhibitor Cocktail (Thermo Fisher Scientific). Protein concentrations of the samples were measured using a BCA Protein Assay kit (Thermo Fisher Scientific). An equal amount of protein (25 μg) was loaded into each lane, separated on 4%–12% Bis-Tris NuPAGE gels (Invitrogen, Carlsbad, CA, USA) and transferred to PVDF membranes (MilliporeSigma) using an XCell SureLock mini-cell system (Invitrogen). After transfer, membranes were blocked with Li-COR Odyssey Blocking buffer (Li-COR Biosciences). The membranes were subsequently probed with the following primary antibodies. Gamma-tubulin was used as an internal control. Anti-Htt (clone mEM48, MAB5374) and anti-γ-tubulin (T6557) antibodies were obtained from Sigma-Aldrich. Anti-GCH-1 (A305-296A-M-1) was obtained from Bethyl Laboratories. Anti-DHFR (ab124814)), anti-TPH2 (ab184505), anti-QDPR/DHPR polyclonal (ab126150), and anti-MTHFR (ab203786) antibodies were obtained from Abcam. Anti-MS (68796), anti-TH (58844S), anti-ChAT (27269) and anti-nNOS (4234) antibodies were purchased from Cell Signaling Technology. Anti-MAT1/2A (NB110-94162) and anti-AHCY (NBP1-55016) antibodies were purchased from Novus Biologicals. Following incubation with primary antibodies, the membranes were probed with IRDye 680RD conjugated to Goat anti-Rabbit IgG or IRDye 800CW conjugated to Goat-anti-Mouse IgG secondary antibody (1:10,000 dilution, LI-COR Biosciences). Image acquisition was performed with a LI-COR Odyssey Infrared Fluorescent scanner (LI-COR Biosciences). All protein bands were quantified using LI-COR software and are expressed as the ratio of each targeted protein band fluorescence intensity to that of γ-tubulin.

### Crystal violet staining

All mice were anesthetized, and the brains were removed and post-fixed in 4% PBS-buffered paraformaldehyde for 24 h. The brain was embedded in paraffin, and a series of consecutive coronal sections (4 μm in thickness) were prepared using the same protocols as described^72^ for crystal violet staining and immunohistochemistry assays. For crystal violet staining, the sections were submerged in graded ethanol and xylene vitrification for demyelination. The sections were stained with 0.1% cresyl violet for 1 min, washed with distilled water, dehydrated with graded ethanol for vitrification, and finally coverslipped. Striatal volumes were calculated using Cavalieri’s principle (volume=*s*_1_*d*_1_+*s*_2_*d*_2_+…+*s*_n_*d*_n_, where *s* is the surface area and *d* is the distance between two sections).

### Immunohistochemistry (IHC) assay

For the immunohistochemistry assay, the prepared sections were submerged in citrate buffer (pH = 6.0) and heated at boiling temperature in a pressure cooker for 10 min for antigen retrieval. The expression level of Htt, GTPCH and DHFR proteins were examined in each group after incubation with primary antibody overnight and HRP-conjugated secondary antibody incubation at 37°C for 45 min. The reaction was visualized with a Pierce™ DAB Substrate Kit (Thermo Fisher Scientific) followed by hematoxylin staining (Abcam) of the nuclei. The number of positively stained cells was counted in five randomly selected microscopic fields at 400X. The average of stained cells was calculated.

### BH_4_ and BH_2_ measurements

To measure BH_4_ and BH_2_ in the plasma of R6/2 and NCAR mice (n = 4), blood (∼ 1 ml) from each mouse was collected by cardiac puncture into an EDTA coated tube containing 100 μl of 1% freshly prepared DTT stock solution to give a final DTT concentration of 0.1% to protect BH_4_ from oxidation as described by Fekkes and Voskuilen-Kooijman^73^. Collected blood was centrifuged at 2,000 x g for 15 min to isolate plasma within 30 min after collection. Plasma was stored at -80°C. To analyze BH_4_ and BH_2_ in the brain (n = 4), brain tissues were first ground with liquid nitrogen. Subsequent sample preparation and LC-MS/MS analysis to measure BH4 and BH2 levels in both plasma and brain tissues were performed as described by Shen et al.^74^.

### Amino acid measurements

To measure the contents of amino acids in the plasma of R6/2 and NCAR mice (n = 4), blood from each mouse was collected by cardiac puncture into an EDTA coated tube. Collected blood was centrifuged at 2,000 x g for 15 min to isolate plasma within 30 min after collection. Collected plasma was stored at -80 °C until analysis. To analyze the contents of amino acids in the brain (n = 4), the brain was excised after anesthetization. Then brain tissues were ground in liquid nitrogen. For detecting amino acids in plasma and brain tissues, the same protocols for sample preparation and Ultraperformance® Liquid Chromatography (UPLC) analysis as described by Peake et al.^75^ were used.

### Statistical Analysis

For all experiments except proteomic analysis, the obtained results are presented as the average ± SD. Statistical significance was analyzed using One-way ANOVA and Student’s t-tests for pairwise mean comparisons (*p* < 0.05). For proteomic analysis, we calculated fold-changes as log2-ratio between average values of Htt_ex1_Q63 vs. Htt_ex1_Q21 groups and performed Student’s t-test with and without Benjamini-Hochberg correction for multiple hypothesis testing.

### Availability of data, materials and methods

The datasets of quantitative proteomic analysis generated for this study were uploaded to MassIVE, and were assigned the dataset identifier MassIVE MSV000084557, which can be accessed at ftp://MSV000084557@massive.ucsd.edu using password 4981 and will be made publicly available once the paper is published. All other data, materials and methods are available from the corresponding authors upon request.

## Supplementary Information

### Supplementary Text

1. PolyQ63 remodeled root proteome extensively
2. PolyQ63 does not affect root hair initiation
3. DAPs possibly responsible for root growth

**References only in Supplementary Text (76-88)**

### Supplementary Tables

Table S1. Metadata of eight samples and three QC arrangements Table S7. Peroxidases involved in ROS generation

Table S8. Mitochondrial membrane DAPs involved in transport

Table S9. Mitochondrial DAPs involved in the electron transfer chain, amino acid metabolism and TCA cycle

Table S10. DAPs involved in the tubular ER network and root hair elongation Table S11. DAPs involved in ribosome biogenesis and ribosome assembly Table S12. DAPs involved in regulation of glutathione redox status

Table S13. RNA polymerases involved in transcription

Table S14. Serine/threonine-protein kinases and phosphatases involved in cell proliferation and expansion

Table S15. DAPs involved in folate and one carbon metabolism

### Supplementary Figures

Fig. S1. Transformation efficiency and regenerated transgenic plants.

Fig. S2. Transgenic plants, leaves, seed pods and seeds.

Fig. S3. PCR and RT-PCR results of seven transgenic lines per genetic cassette.

Fig. S4. Detection of protein aggregates in plant leaves.

Fig. S5. Principal component analysis of quantified proteins.

Fig. S6. Selected top 10 GO results of abundance-decreased DAPs.

Fig. S7. Selected top 10 GO results of GO results of abundance-increased DAPs.

Fig. S8. ATP levels of young leaf tissues.

Fig. S9. Root hair initiation in induced roots.

Fig. S10. Normal root tip structure and DAPs associated with root cell division, elongation or expansion.

Fig. S11. Characterization of 4-week old wide-type non-carrier (NCAR) and R6/2 HD mice.

Fig. S12. Representative Western blotting results of GTPCH, DHFR, QDPR, MS, MAT1/2A, AHCY, MTHFR, TPH2, TH, nNOS and ChAT.

### Other Supplementary Tables in Excel spreadsheets

Table S2. Identified 44,888 peptides

Table S3. Identified 7,189 proteins

Table S4. 5,073 proteins quantified by ≥ 2 peptides

Table S5. 1,693 DAPs between Htt_ex1_Q63 and Htt_ex1_Q21

Table S6. GO enriched pathways

## Acknowledgments

We thank M.W. Foster and M.A. Moseley for proteomic analysis, V.M. Knowlton for the help with TEM, W.Z. Duan for discussions and critical reading of the manuscript, Y.-H. Sun for helpful discussions in GO analysis, and S.H. Li for providing PCR buffer information. This work was supported by grants from the National Institute of General Medical Sciences (SC1GM111178) to J.X. and a scholarship from the China Scholarship Council (201306305041) to C.Z.

## Conflict of interest

The authors declare no conflict of interest.

## Author contributions

T.D. and J.X. designed research; C.-Y.H, C.S., F.S.K., M.T.H., E.A., G.S.J., M.D.T., T.D., and J.X. performed the experiments; C.-Y.H, F.S.K., E.A., T.D. and J.X. analyzed the data. C.-Y.H, F.S.K. and J.X. drafted the manuscript with input from all authors.

## Notes

### Competing Interest Statement

The authors have declared no competing interest.

## References

1. Gusella, J. F. & MacDonald, M. E. Molecular genetics: unmasking polyglutamine triggers in neurodegenerative disease. Nat. Rev. Neurosci. 1, 109–115 (2000).

2. Di Prospero, N. A. & Fischbeck, K. H. (2005) Therapeutics development for triplet repeat expansion diseases. Nat. Rev. Genet. 6, 756–765 (2005).

3. The Huntington’s Disease Collaborative Research Group. A novel gene containing a trinucleotide repeat that is expanded and unstable on Huntington’s disease chromosomes. Cell 72, 971–983 (1993).

4. Veldman, M. B. & Yang, X. W. Molecular insights into cortico-striatal miscommunications in Huntington’s disease. Curr. Opin. Neurobiol. 48, 79–89 (2018).

5. Genetic Modifiers of Huntington’s Disease (GeM-HD) Consortium. CAG repeat not polyglutamine length determines timing of Huntington’s disease onset. Cell 178, 887-900.e14 (2019).

6. Wertz, M. H. et al. (2020) Genome-wide in vivo CNS screening identifies genes that modify CNS neuronal survival and mHTT toxicity. Neuron 106, 76–89.e8 (2020).

7. Brenner, E.D. et al. Plant neurobiology: an integrated view of plant signaling. Trends Plant Sci. 11, 413–419 (2006).

8. Baluska, F. Recent surprising similarities between plant cells and neurons. Plant Signal. Behav. 5, 87–89 (2010).

9. Guttman, D. C. Plants as models for the study of human pathogenesis. Biotechnol. Adv. 22, 363–382 (2004).

10. Spampinato, C. P. & Gomez-Casati, D. F. Research on plants for understanding of disease of nuclear and mitochondrial origin. J. Biomed. Biotechnol. 2012, 836196 (2012).

11. Hanson, A. D., Gage, D. A. & Shachar-Hill, Y. Plant one-carbon metabolism and its engineering. Trends Plant Sci. 5, 206–213 (2000).

12. Gorelova, V., Ambach, L., Rébeillé, F., Stove, C. & Van Der Straeten, D. Folates in Plants: Research advances and progress in crop biofortification. Front. Chem. 5, 21 (2017).

13. Coppedè, F. One-carbon metabolism and Alzheimer’s disease: focus on epigenetics. Curr. Genomics 11, 246–260 (2010).

14. Thomas, E. A. DNA methylation in Huntington’s disease: Implications for transgenerational effects. Neurosci. Lett. 625, 34–39 (2016).

15. Ducker, G. S. & Rabinowitz, J. D. One-carbon metabolism in health and disease. Cell Metab. 25, 27–42 (2017).

16. Mohd Murshid, N., Aminullah Lubis, F. & Makpol, S. Epigenetic changes and its intervention in age-related neurodegenerative diseases. Cell. Mol. Neurobiol. 42, 577–595 (2022).

17. Kapatos G. The neurobiology of tetrahydrobiopterin biosynthesis: a model for regulation of GTP cyclohydrolase I gene transcription within nigrostriatal dopamine neurons. IUBMB Life 65, 323–333 (2013).

18. Pouladi, M. A., Morton, A. J. & Hayden, M. R. Choosing an animal model for the study of Huntington’s disease. Nat. Rev. Neurosci. 14, 708–721 (2013).

19. Chang, R., Liu, X., Li, S. H. & Li, X. J. Transgenic animal models for study of the pathogenesis of Huntington’s disease and therapy. Drug Des. Devel. Ther. 9, 2179–2188 (2015).

20. Li, S. H. & Li, X. J. Aggregation of *N*-terminal huntingtin is dependent on the length of Its glutamine repeats. Hum. Mol. Genet. 7, 777–782 (1998).

21. Vonsattel, J. P. & DiFiglia, M. Huntington disease. J. Neuropathol. Exp. Neurol. 57, 369–384 (1998).

22. Landles, C. & Bates, G. P. Huntingtin and the molecular pathogenesis of Huntington’s disease. Fourth in molecular medicine review series. EMBO Rep. 5, 958–963 (2004).

23. Kim, Y. E. et al. Soluble oligomers of polyQ-expanded huntingtin target a multiplicity of key cellular factors. Mol. Cell 63, 951–964 (2017).

24. Hu, J. et al. A class of dynamin-like GTPases involved in the generation of the tubular ER network. Cell 138, 549–561 (2009).

25. Chen, J., Stefano, G., Brandizzi, F. & Zheng, H. Arabidopsis RHD3 mediates the generation of the tubular ER network and is required for Golgi distribution and motility in plant cells. J. Cell. Sci. 124, 2241–2252 (2011).

26. Lee, H. et al. An Arabidopsis reticulon and the atlastin homologue RHD3-like2 act together in shaping the tubular endoplasmic reticulum. New Phytol. 197, 481–489 (2013).

27. Oláh, J. et al. Increased glucose metabolism and ATP level in brain tissue of Huntington’s disease transgenic mice. FEBS J. 275, 4740–4755 (2008).

28. Bossy-Wetzel, E., Petrilli, A. & Knott, A. B. Mutant huntingtin and mitochondrial dysfunction. Trends Neurosci. 31, 609–616 (2008).

29. Hosp, F. et al. Spatiotemporal proteomic profiling of Huntington’s disease inclusions reveals widespread loss of protein function. Cell Rep. 21, 2291–2303 (2017).

30. Libault, M., Brechenmacher, L., Cheng, J., Xu, D. & Stacey, G. Root hair systems biology. Trends Plant Sci.15, 641–650 (2010).

31. Balcerowicz, D., Schoenaers, S. & Vissenberg, K. (2015). Cell fate determination and the switch from diffuse growth to planar polarity in Arabidopsis root epidermal cells. Front. Plant Sci. 6, 1163 (2015).

32. Mencacci, N. E. et al. Parkinson’s disease in GTP cyclohydrolase 1 mutation carriers. Brain 137, 2480–2492 (2014).

33. Raychaudhuri, S., Sinha, M., Mukhopadhyay, D. & Bhattacharyya, N. P. HYPK, a Huntingtin interacting protein, reduces aggregates and apoptosis induced by *N*-terminal Huntingtin with 40 glutamines in Neuro2a cells and exhibits chaperone-like activity. Hum. Mol. Genet. 17, 240–255 (2008).

34. Wang, H., Lockwood, S. K., Hoeltzel, M. F. & Schiedelbein, J. W. The ROOT HAIR DEFECTIVE3 gene encodes an evolutionarily conserved protein with GTP-binding motifs and is required for regulated cell enlargement in Arabidopsis. Genes Dev. 11, 799–811 (1997).

35. Aberg, K., Saetre, P., Jareborg, N. & Jazin, E. Human QKI, a potential regulator of mRNA expression of human oligodendrocyte-related genes involved in schizophrenia. Proc. Natl. Acad. Sci. USA 103, 7482–7487 (2006).

36. Forde, B. G. & Lea, P. J. Glutamate in plants: metabolism, regulation, and signaling. J. Exp. Bot. 58, 2339–2358 (2007).

37. Basset, G. et al. Folate synthesis in plants: the first step of the pterin branch is mediated by a unique bimodular GTP cyclohydrolase I. Proc. Natl. Acad. Sci. USA 99, 12489–12494 (2002).

38. Hersch, S. M. & Ferrante, R. J. (2004) Translating therapies for Huntington’s disease from genetic animal models to clinical trials. NeuroRx 1, 298–306 (2004).

39. Kurian, M. A., Gissen, P., Smith, M., Heales, S. Jr. & Clayton, P. T. The monoamine neurotransmitter disorders: an expanding range of neurological syndromes. Lancet Neurol. 10, 721–733 (2011).

40. Xu, F. et al. Disturbed biopterin and folate metabolism in the Qdpr-deficient mouse. FEBS Lett. 588, 3924–3931 (2014).

41. Luo, M., Piffanelli, P., Rastelli, L. & Cella, R. Molecular cloning and analysis of a cDNA coding for the bifunctional dihydrofolate reductase-thymidylate synthase of *Daucus carota*. Plant Mol. Biol. 22, 427–435 (1993).

42. Rangel-Barajas, C. & Rebec, G.V. Dysregulation of corticostriatal connectivity in Huntington’s Disease: A role for dopamine modulation. J. Huntingtons Dis. 5, 303–331 (2016).

43. Fanet, H. et al. Tetrahydrobiopterin administration facilitates amphetamine-induced dopamine release and motivation in mice. Behav. Brain Res. 379, 112348 (2020).

44. Newman, A. C. & Maddocks, O. D. K. One-carbon metabolism in cancer. Br. J. Cancer 116, 1499–1504 (2017).

45. Reina-Campos, M., Diaz-Meco, M. T. & Moscat, J. The complexity of the serine glycine one-carbon pathway in cancer. J. Cell Biol. 219, e201907022 (2020).

46. Berry, M. D. Mammalian central nervous system trace amines. Pharmacologic amphetamines, physiologic neuromodulators. J. Neurochem. 90, 257–271 (2004).

47. Pei, Y., Asif-Malik, A. & Canales, J. J. Trace amines and the trace amine-associated receptor 1: pharmacology, neurochemistry, and clinical implications”. Front. Neurosci. 10, 148 (2016).

48. Albrecht, J., Sidoryk-Węgrzynowicz, M., Zielińska, M. & Aschner, M. Roles of glutamine in neurotransmission. Neuron Glia Biol. 6, 263–276 (2011).

49. Zielińska, M., Ruszkiewicz, J., Hilgier, W., Fręśko, I. & Albrecht, J. Hyperammonemia increases the expression and activity of the glutamine/arginine transporter y+ LAT2 in rat cerebral cortex: implications for the nitric oxide/cGMP pathway. Neurochem. Int. 58, 190–195 (2011).

50. Emery, P. W. Amino acids: Metabolism. In Encyclopedia of Human Nutrition 3rd edn (ed. Caballero, B.), Academic Press, 72–78 (2013).

51. Walters, D. C. et al. Metabolomic analyses of vigabatrin (VGB)-treated mice: GABA- transaminase inhibition significantly alters amino acid profiles in murine neural and non- neural tissues. Neurochem. Int. 125, 151–162 (2019).

52. Mochel, F. et al. Early energy deficit in Huntington disease: identification of a plasma biomarker traceable during disease progression. PLoS One 2, e647 (2007).

53. Mochel, F., Benaich, S., Rabier, D. & Durr, A. Validation of plasma branched chain amino acids as biomarkers in Huntington disease. Arch. Neurol. 68, 265–267 (2011).

54. Skene, D. J. et al. Metabolic profiling of presymptomatic Huntington’s disease sheep reveals novel biomarkers. Sci. Rep. 7, 43030 (2017).

55. Wu, G. Amino acids: metabolism, functions, and nutrition. Amino acids 37: 1–17 (2009).

56. Moss, D. J. H. et al. Identification of genetic variants associated with Huntington’s disease progression: a genome-wide association study. Lancet Neurol. 16, 701–711 (2017).

57. Jiang, X. et al. A novel GTPCH deficiency mouse model exhibiting tetrahydrobiopterin- related metabolic disturbance and infancy-onset motor impairments. Metabolism 94, 96–104 (2019).

58. Glajch, K. E. & Sadri-Vakili, G. Epigenetic mechanisms involved in Huntington’s disease pathogenesis. J. Huntingtons Dis. 4, 1–15 (2015).

59. Sharma, S. & Taliyan, R. Transcriptional dysregulation in Huntington’s disease: The role of histone deacetylases. Pharmacol. Res. 100, 157–169 (2015).

60. Raymond, L. A. et al. Pathophysiology of Huntington’s disease: time-dependent alterations in synaptic and receptor function. Neuroscience 198, 252–273 (2011).

61. Moreno-Jiménez, E.P. et al. Adult hippocampal neurogenesis is abundant in neurologically healthy subjects and drops sharply in patients with Alzheimer’s disease. Nat. Med. 25, 554–560 (2019).

62. Musa, T. A., Hung, C.-Y., Darlington, D. E., Sane, D. C. & Xie, J. H. Overexpression of human erythropoietin in tobacco does not affect plant fertility or morphology. Plant Biotechnol. Rep. 3, 157–165 (2009).

63. Holsters, M. et al. Transfection and transformation of *Agrobacterium tumefaciens*. Mol. Gen. Genet. 163, 181–187 (1978).

64. Twyman, R. M., Stoger, E., Schillberg, S., Christou, P. & Fischer, R. Molecular farming in plants: host systems and expression technology. Trends Biotechnol. 21, 570–578 (2003).

65. Dewey, R. E. & Xie, J. H. Molecular genetics of alkaloid biosynthesis in *Nicotiana tabacum*. Phytochemistry 94, 10–27 (2013).

66. Horsch, R. B. et al. *Leaf disc transformation* in *Plant molecular biology manual* (Gelvin, S. B. & Schilperoot, R. A. Eds) (Kluwer Academic), pp. 1–9 (1988).

67. Jama, M., Millson, A., Miller, C. E. & Lyon, E. Triplet repeat primed PCR simplifies testing for Huntington disease. J. Mol. Diagn. 15, 255–262 (2013).

68. Huang, B. et al. Mutant huntingtin down-regulates Myelin Regulatory Factor-mediated myelin gene expression and affects mature oligodendrocytes. Neuron 85, 1212–1226 (2015).

69. Tian, T. et al. AgriGO v2.0: a GO analysis toolkit for the agricultural community, 2017 update. Nucleic Acids Res. 45(W), W122–W129 (2017).

70. Deng, W., Wang, Y., Liu, Z., Cheng, H. & Xue, Y. HemI: a toolkit for illustrating heatmaps. PLoS One. 9, e111988 (2014).

71. Babu, M. M. *An introduction to microarray data analysis* in *Computational Genomics: Theory and Application* (Grant, R. P. Ed) (Horizon Bioscience), pp. 225–249 (2004).

72. Mehta, S. L., Kumari, S., Mendelev, N. & Li, P. A. (2012) Selenium preserves mitochondrial function, stimulates mitochondrial biogenesis, and reduces infarct volume after focal cerebral ischemia. BMC Neurosci. 13, 79 (2012).

73. Fekkes, D., Voskuilen-Kooijman, A. Quantitation of total biopterin and tetrahydrobiopterin in plasma. Clin. Biochem. 40, 411–413 (2007).

74. Shen, J. S. et al. Tetrahydrobiopterin deficiency in the pathogenesis of Fabry disease. Hum. Mol. Genet. 26, 1182–1192 (2017).

75. Peake, R. W. A. et al. Improved separation and analysis of plasma amino acids by modification of the MassTrak™ AAA Solution Ultraperformance® liquid chromatography method. Clin. Chim. Acta 423, 75–82 (2013).

