## Supplemental tables and figures for "Mutant huntingtin exon-1 impaired GTPCH and DHFR expression in plants and mice"

**This PDF file includes:**

**Supplementary Text**

1. PolyQ63 remodeled root proteome extensively
2. PolyQ63 does not affect root hair initiation
3. DAPs possibly responsible for root growth

**References only in Supplementary Text (76-88)**

**Supplementary Tables**

Table S1. Metadata of eight samples and three QC arrangements

Table S7. Peroxidases involved in ROS generation

Table S8. Mitochondrial membrane DAPs involved in transport

Table S9. Mitochondrial DAPs involved in the electron transfer chain, amino acid metabolism and TCA cycle

Table S10. DAPs involved in the tubular ER network and root hair elongation

Table S11. DAPs involved in ribosome biogenesis and ribosome assembly

Table S12. DAPs involved in regulation of glutathione redox status

Table S13. RNA polymerases involved in transcription

Table S14. Serine/threonine-protein kinases and phosphatases involved in cell proliferation and expansion

Table S15. DAPs involved in folate and one carbon metabolism

**Supplementary Figures**

Fig. S1. Transformation efficiency and regenerated transgenic plants.

Fig. S2. Transgenic plants, leaves, seed pods and seeds.

Fig. S3. PCR and RT-PCR results of seven transgenic lines per genetic cassette.

Fig. S4. Detection of protein aggregates in plant leaves.

Fig. S5. Principal component analysis of quantified proteins.

Fig. S6. Selected top 10 GO results of abundance-decreased DAPs.

Fig. S7. Selected top 10 GO results of GO results of abundance-increased DAPs.

Fig. S8. ATP levels of young leaf tissues.

Fig. S9. Root hair initiation in induced roots.

Fig. S10. Normal root tip structure and DAPs associated with root cell division, elongation or expansion.

Fig. S11. Characterization of 4-week old wide-type non-carrier (NCAR) and R6/2 HD mice.

Fig. S12. Representative Western blotting results of GTPCH, DHFR, QDPR, MS,

MAT1/2A, AHCY, MTHFR, TPH2, TH, nNOS and ChAT.

**Other Supplementary Tables in Excel spreadsheets:**

Table S2. Identified 44,888 peptides

Table S3. Identified 7,189 proteins

Table S4. 5,073 proteins quantified by ≥ 2 peptides

Table S5. 1,693 DAPs between $Htt_{ex1}$Q63 and $Htt_{ex1}$Q21

Table S6. GO enriched pathways

### Supplementary Text

#### 1. PolyQ63 remodeled root proteome extensively

To get insight into polyQ63-mediated changes in root proteome, quantitative proteomic analysis was performed with $Htt_{ex1}$Q63 and $Htt_{ex1}$Q21 young roots. A total of 44,888 peptides (Table S2) and 7,189 proteins (Table S3) were quantified across all samples (database: ftp://MSV000084557@massive.ucsd.edu) after removing low quality peptides with poor chromatographic peak shape, and scaling of the data to the robust mean across all samples. To determine the quality of quantitative proteomic analysis, both analytical variability and biological variability were analyzed. To assess technical reproducibility, the percentage (%) of coefficient of variation (CV) was calculated for each protein across the three injections of a QC pool that were interspersed throughout the study while %CVs were also measured for each protein across the individual analyses to assess biological variability. The mean %CV of the QC pools was 13.9% for all proteins and 11.5% for proteins quantified by 2 or more peptides (Table S3). Variability of the biological samples (analytical variability plus biological variability) was

25.6% for proteins or 22.9% for proteins quantified by 2 or more peptides (Table S3). The results of %CV indicate that the technical reproducibility was good, which was much higher than the observed biological reproducibility. These data were reflected in the clustering of the QC samples as visualized by PCA analysis (Fig. S5) while the individual samples clustered based on treatment group. Therefore, these detected proteins were used for their differential expression analysis.

Of these 7,189 proteins, 5,073 of them were further quantified by 2 or more peptides (Table S4). A close examination revealed that abundance of 2,696 proteins was decreased while that of 2,377 proteins was increased (Table S4). When a criterion for higher confidence of identification and quantification with a CV<30% across QC pool replicates, a FDR-corrected $p<0.05$, and the cutoff of ≥1.5-fold changes in abundance was used, a total of 1,693 DAPs were identified between $Htt_{ex1}Q63$ and $Htt_{ex1}Q21$ roots (Table S5). Among them, there were 854 abundance-increased and 839 abundance-decreased DAPs in $Htt_{ex1}Q63$ compared to $Htt_{ex1}Q21$. GO enrichment analysis of these DAPs showed that polyQ63 primarily reduced abundances of proteins involved in translation (GO:0006412, $p$ = 2.94e-06; GO:0000462, $p$ = 1.26e-11), folding (GO:0006457, $p$ = 0.000396), and transport (GO:0015031, $p$ = 4.86e-09), especially from vesicle-mediated ER to Golgi (GO:0006888, $p$=2.89e-06) (Fig. S6; Table S6). The impairment of protein transport and localization is further supported by affected proteins involved in vesicle (GO:0031988, $p$=0.000321), ER (GO:0005789, $p$=7.6e-12), Golgi (GO:0000139, $p$ = 1.91e-06), inner mitochondrial membrane (GO:0098800, $p$=0.0386), and macromolecular complexes (GO:0032991, $p$=4.03e-14). When DAPs were categorized based on molecular functions, polyQ63 mainly interrupted the abundances of proteins having activities in the transfer of C1 units (GO:0016741, $p$=4.22e-07), GTPase binding (GO:0051020, $p$=0.000325), RNA binding

(GO:0003723, *p*=1.6e-13), structural constituent of ribosome (GO:0003735; *p*=1e-07) and unfolded protein binding (GO:0051082 *p*=5.74e-06).

Among abundance-increased DAPs, the majority were oxidative stress-related proteins involved in producing reactive oxygen species (ROS) and responding to high ROS levels (GO:0006979, *p*=1.65e-11; GO:0072593, *p*=4.53e-11), as well as enzymes involved in various metabolic processes, such as the carbohydrate metabolic process (GO:0005975, *p*=4.3e-09), hydrogen peroxide catabolic process (GO:0042744, *p*=9.99e-12), and carboxylic acid metabolic process (GO:0019752, *p*=3.71e-06) (Fig. S7; Table S6). Proteins localized at extracellular region (GO:0005576, *p*=1.28e-11) were also significantly enriched. In addition, mitochondrial membrane proteins were enriched (GO:0031966, *p*=0.00464). In term of molecular function, some abundance-increased proteins were enriched in regulating the activities of hydrolases (GO:0016787, *p*=3.42e-07), glutathione transferases (GO:0004364, *p*=4.44e-09), transaminases (GO:0008483, *p*=4.15e-06), oxidoreductases (GO:0016491, *p*=1.06e-26), peroxidases (GO:0004601, *p*=9.66e-14), lyases (GO:0016829, *p*=4.67e-06) as well as proteins with binding activities of ions (GO:0043167, *p*=1e-05), cofactors (GO:0048037, *p*=1.5e-11) and heme (GO:0020037, *p*=3.73e-08). The analysis of these identified DAPs demonstrated that polyQ63 caused widespread remodeling of root proteome.

### 2. PolyQ63 does not affect root hair initiation

In *Arabidopsis*, root hair initiation begins with the establishment of polarity within epidermal cells[76,77] and whether an epidermal cell can be differentiated into root hair (H) or non-hair (N) cell depends on the absence or presence of an inhibitory pathway consisting of the following transcription factors GLABRA2 (GL2), CAPRICE (CPC), TRANSPARENT TESTA GLABRA

(TTG), GLABRA3 (GL3)/ENHANCER OF GLABRA3 (EGL3) and WEREWOLF (WER) (Fig. S9A)[30]. An epidermal cell becomes H cell when CPC-TTG-GL3-EGL3 complex is formed to inactivate an inhibitory transcription factor GL2 of the final step of H cell formation whereas a cell becomes N cell when WER-TTG-GL3-EGL3 complex is formed to promote GL2 activity[30]. To understand whether the phenomenon of no or less root hairs in induced adventitious roots from $Htt_{ex1}$Q63 shoots is due to affected root hair initiation or outgrowth, we examined these transcription factors related to root hair initiation in our proteomic data (Table S4). Among them, GL2 was found to be reduced 1.5-fold in $Htt_{ex1}$Q63 roots compared to $Htt_{ex1}$Q21 (Fig. S9B; Table S4). Although none of other transcription factors were detected among the DAPs, the decreased abundance of GL2 would benefit or at least not inhibit H cell initiation in $Htt_{ex1}$Q63 roots, suggesting that polyQ63 might not affect root hair initiation at this stage. Indeed, the adventitious roots were induced having hairs when $Htt_{ex1}$Q63 shoots were inoculated on the surface of solid maintenance medium (Fig. S9C).

**3. DAPs possibly responsible for root growth**

A root tip consists of meristematic, transition, elongation and maturation zones based on their characteristic cellular activities (Fig. S10A)[31,78]. Our proteomic data suggest that the following DAPs are possibly responsible for the observed restricted root growth and unorganized root cap structure in $Htt_{ex1}$Q63 (Fig. S10B and S10C). Firstly, nine DAPs associated with cell division, expansion and elongation[34,79-84] were abundance-decreased (Fig. S10B). Secondly, eight ribosome biogenesis proteins together with a group of ribosome assembly factors, including ATPases, GTPases, helicases, and kinases were also among the abundance-decreased DAPs (Fig. S10C-I; Table S11). Ribosome biogenesis is tightly regulated and inextricably linked to cell division and growth[85]. Impairment of ribosome biogenesis by mHtt has also been observed in an animal HD model[86] and a N2a cell HD model[23]. Thirdly, as many as 27 glutathione *S*-transferases and 10 glutathione metabolism-related proteins were abundance-decreased DAPs (Fig. S10C-II; Table S12), suggesting that glutathione redox status was altered. In plants, it is also known that altered glutathione redox status does not favor either root growth or root apical meristem maintenance[87]. Fourthly, all seven DNA-directed RNA polymerases were found to be abundance-decreased (Fig. S10C-III; Table S13), as also observed in an animal HD model[86]. Lastly, 11 serine/threonine-protein kinases and phosphatases were among abundance-decreased DAPs (Fig. S10C-IV; Table S14), which are known to be involved in cell proliferation and expansion[88]. Since many of these abundance-decreased DAPs play important roles in root growth and development, they may be responsible for defective root growth and root cap integrity.

**References only in Supplementary Materials (76-88):**


76. Carol, R. J. & Dolan, L. Building a hair: tip growth in *Arabidopsis thaliana* root hairs. *Philos. Trans. R. Soc. Lond. B Biol. Sci.* **357**, 815-821 (2002).
77. Carol, R. J. & Dolan, L. The role of reactive oxygen species in cell growth: lessons from root hairs. *J. Exp. Bot.* **57**, 1829-1834 (2006).
78. Anderson, C. T., Carroll, A., Akhmetova, L. & Somerville, C. Real-time imaging of cellulose reorientation during cell wall expansion in Arabidopsis roots. *Plant Physiol.* **152**, 787-796 (2010).
79. Burk, D. H., Liu, B., Zhong, R., Morrison, W. H. & Ye, Z. H. A katanin-like protein regulates normal cell wall biosynthesis and cell elongation. *Plant Cell* **13**, 807-827 (2001).
80. Wagner, T. A. & Kohorn, B. D. Wall-associated kinases are expressed throughout plant development and are required for cell expansion. *Plant Cell* **13**, 303-318 (2001).

81. Burger, C., Rondet, S., Benveniste, P. & Schaller, H. Virus-induced silencing of sterol biosynthetic genes: identification of a *Nicotiana tabacum* L. obtusifoliol-14α-demethylase (CYP51) by genetic manipulation of the sterol biosynthetic pathway in *Nicotiana benthamiana* L. *J. Exp. Bot.* **54**, 1675-1683 (2003).
82. Soyano, T., Nishihama, R., Morikiyo, K., Ishikawa, M. & Machida, Y. NQK1/NtMEK1 is a MAPKK that acts in the NPK1 MAPKKK-mediated MAPK cascade and is required for plant cytokinesis. *Genes Dev.* **17**, 1055-1067 (2003).
83. Brown, D. M., Zhang, Z., Stephens, E., Dupree, P. & Turner, S. R. Characterization of IRX10 and IRX10-like reveals an essential role in glucuronoxylan biosynthesis in Arabidopsis. *Plant J.* **57**, 732-746 (2009).
84. Hermans, C., Porco, S., Verbruggen, N. & Bush, D. R. Chitinase-like protein CTL1 plays a role in altering root system architecture in response to multiple environmental conditions. *Plant Physiol.* **152**, 904-917 (2010).
85. Thomson, E., Ferreira-Cerca, S. & Hurt, E. Eukaryotic ribosome biogenesis at a glance. *J. Cell Sci.* **126**, 4815-4821 (2013).
86. Shirasaki, D. I. et al. Network organization of the huntingtin proteomic interactome in mammalian brain. *Neuron* **75**, 41-57 (2012).
87. Yu, X. et al. Plastid-localized glutathione reductase2-regulated glutathione redox status is essential for Arabidopsis root apical meristem maintenance. *Plant Cell* **25**, 4451-4468 (2013).
88. Uhrig, R. G., Labandera, A. M. & Moorhead, G. B. Arabidopsis PPP family of serine/threonine protein phosphatases: many targets but few engines. *Trends Plant Sci.* **18**, 505-513 (2013).

**Table S1. Metadata of eight samples and three QC arrangements**

| Samples | File Name | Run order |
|---|---|---|
| QC1 | ID38360_04_06069L_4981_052318 | 1 |
| $Htt_{ex1}$Q21-1 | ID38304_01_06069L_4981_052318 | 2 |
| $Htt_{ex1}$Q63-2 | ID38309_01_06069L_4981_052318 | 3 |
| $Htt_{ex1}$Q21-3 | ID38306_01_06069L_4981_052318 | 4 |
| $Htt_{ex1}$Q63-3 | ID38310_01_06069L_4981_052318 | 5 |
| QC2 | ID38360_05_06069L_4981_052318 | 6 |
| $Htt_{ex1}$Q63-1 | ID38308_01_06069L_4981_052318 | 7 |
| $Htt_{ex1}$Q21-2 | ID38305_01_06069L_4981_052318 | 8 |
| $Htt_{ex1}$Q63-4 | ID38311_01_06069L_4981_052318 | 9 |
| $Htt_{ex1}$Q21-4 | ID38307_01_06069L_4981_052318 | 10 |
| QC3 | ID38360_06_06069L_4981_052318 | 11 |

**Table S7. Peroxidases involved in ROS generation**

| UniProtKB | Protein description | Log2 fold-change | *p*-value* |
|---|---|---|---|
| A0A1S3YET7_TOBAC | Peroxidase | -1.64 | 9.4E-04 |
| A0A1S4DNP1_TOBAC | Peroxidase | -1.41 | 6.3E-05 |
| A0A1S3ZXH0_TOBAC | Peroxidase | -0.81 | 1.0E-03 |
| A0A1S3XNL8_TOBAC | Peroxidase | 2.56 | 5.4E-05 |
| A0A1S4BNU3_TOBAC | Peroxidase | 2.51 | 5.8E-05 |
| PERN1_TOBAC | Peroxidase N1 | 2.42 | 2.2E-05 |
| A0A1S4D2U8_TOBAC | Peroxidase | 2.18 | 5.3E-04 |
| A0A1S4CK25_TOBAC | Peroxidase | 2.16 | 5.6E-06 |
| A0A1S4AW97_TOBAC | Peroxidase | 2.12 | 7.5E-05 |
| A0A1S3Z5I5_TOBAC | L-ascorbate peroxidase 3, peroxisomal-like | 1.91 | 1.3E-02 |
| A0A1S4A7R5_TOBAC | Peroxidase | 1.88 | 2.5E-04 |
| A0A1S4C127_TOBAC | Peroxidase | 1.83 | 3.0E-04 |
| A0A1S3WXW1_TOBAC | Peroxidase | 1.80 | 4.3E-05 |
| Q50LG5_TOBAC | Peroxidase | 1.39 | 1.0E-03 |
| A0A1S3YS19_TOBAC | Peroxidase | 1.38 | 2.0E-03 |
| A0A1S3YKK2_TOBAC | Peroxidase | 1.31 | 7.0E-03 |
| Q40487_TOBAC | Peroxidase | 1.27 | 1.0E-04 |
| A0A1S4BJB5_TOBAC | Peroxidase | 1.18 | 1.3E-04 |
| A0A1S4BEZ8_TOBAC | Peroxidase | 1.13 | 6.8E-04 |
| A0A1S4D627_TOBAC | Peroxidase | 1.10 | 1.0E-03 |
| A0A1S3Z4R4_TOBAC | Peroxidase | 1.08 | 3.6E-05 |
| A0A1S3XNR7_TOBAC | Glutathione peroxidase | 1.02 | 6.2E-05 |

| | | | |
|---|---|---|---|
| A0A1S4CKJ4_TOBAC | L-ascorbate peroxidase 3, peroxisomal | 1.02 | 7.5E-06 |
| A0A1S4CXA3_TOBAC | Peroxidase | 1.00 | 4.9E-05 |
| A0A1S3XQB7_TOBAC | Peroxidase | 0.88 | 1.7E-02 |
| A0A1S3XBP1_TOBAC | Peroxidase | 0.87 | 2.0E-03 |
| A0A1S4C0F0_TOBAC | Peroxidase | 0.87 | 6.4E-05 |
| A0A1S4AIS4_TOBAC | Peroxidase | 0.75 | 1.1E-02 |
| A0A1S3YB42_TOBAC | Glutathione peroxidase | 0.65 | 4.8E-04 |
| A0A1S3ZT94_TOBAC | Peroxidase | 0.63 | 8.0E-03 |

*: t-test with FDR correction

**Table S8. Mitochondrial membrane DAPs involved in transport**

| UniProtKB | Protein description | Log2 fold-change | *p*-value* |
|---|---|---|---|
| A0A1S3ZIV3_TOBAC | Mitochondrial import inner membrane translocase subunit Tim9 | -1.51 | 1.0E-03 |
| A0A1S4CCA3_TOBAC | Mitochondrial import inner membrane translocase subunit Tim13-like | -1.37 | 3.3E-04 |
| A0A1S4A7X7_TOBAC | Mitochondrial inner membrane protein OXA1-like | -1.31 | 5.4E-04 |
| A0A1S3YGL2_TOBAC | Probable mitochondrial import inner membrane translocase subunit TIM21 | -0.77 | 8.4E-04 |
| A0A1S4AKN6_TOBAC | Mitochondrial inner membrane protease ATP23 | -0.76 | 2.4E-02 |
| A9CM21_TOBAC | Voltage-dependent anion channel | 0.92 | 6.1E-05 |
| A0A1S4CVQ8_TOBAC | LETM1 and EF-hand domain-containing protein 1, mitochondrial-like | 0.86 | 2.6E-05 |
| A0A1S4CBH2_TOBAC | Mitochondrial outer membrane protein porin 2-like | 0.84 | 1.4E-04 |
| A0A1S3XKP1_TOBAC | Mitochondrial pyruvate carrier | 0.83 | 4.6E-05 |
| A0A1S4B0F0_TOBAC | Mitochondrial pyruvate carrier | 0.80 | 1.2E-04 |
| A0A1S3YFU4_TOBAC | LETM1 and EF-hand domain-containing protein 1, mitochondrial-like | 0.77 | 1.8E-05 |
| A0A1S3ZVC1_TOBAC | Mitochondrial pyruvate carrier | 0.76 | 8.7E-05 |
| A0A1S3XCA7_TOBAC | Mitochondrial uncoupling protein 5-like | 0.76 | 3.0E-03 |
| A0A1S4DJZ0_TOBAC | Mechanosensitive ion channel protein 1, mitochondrial-like isoform X1 | 0.62 | 4.0E-03 |

*: t-test with FDR correction

**Table S9. Mitochondrial DAPs involved in electron transfer chain, amino acid metabolism and TCA cycle**

| UniProtKB | Protein description | Log2 fold-change | *p*-value* |
|---|---|---|---|
| **Electron transfer chain** | | | |
| A0A1S4CJ56_TOBAC | Cytochrome c oxidase assembly protein COX11, mitochondrial-like isoform X1 | -0.79 | 1.2E-02 |
| A0A1S4CBE2_TOBAC | Mitochondrial zinc maintenance protein 1, mitochondrial-like | -0.64 | 8.5E-04 |
| A0A1S4D561_TOBAC | 2-Methylacyl-CoA dehydrogenase, mitochondrial-like isoform X1 | 2.30 | 7.5E-06 |
| A0A1S3ZHK0_TOBAC | Formate dehydrogenase, mitochondrial | 1.92 | 3.9E-04 |
| A0A1S3Y6A0_TOBAC | Electron transfer flavoprotein-ubiquinone oxidoreductase, mitochondrial | 1.34 | 2.4E-05 |
| A0A1S3XVJ3_TOBAC | Uncharacterized protein At2g27730, mitochondrial-like isoform X1 | 1.31 | 3.2E-05 |
| A0A1S3XC77_TOBAC | Electron transfer flavoprotein subunit alpha, mitochondrial-like | 1.23 | 3.0E-05 |
| AOX1_TOBAC | Ubiquinol oxidase 1, mitochondrial | 1.15 | 9.8E-05 |
| A0A1S3ZC77_TOBAC | Electron transfer flavoprotein subunit beta, mitochondrial-like | 1.14 | 3.5E-05 |
| A0A1S3ZUW5_TOBAC | Internal alternative NAD(P)H-ubiquinone oxidoreductase A1, mitochondrial-like | 1.04 | 1.6E-05 |
| A0A1S4DIL6_TOBAC | Uncharacterized protein At2g27730, mitochondrial-like | 0.89 | 6.0E-05 |
| A0A1S3Z2K5_TOBAC | Aldehyde dehydrogenase family 2 member B7, mitochondrial-like | 0.85 | 3.8E-05 |
| A0A1S4CQH8_TOBAC | Ubiquinone biosynthesis monooxygenase COQ6, mitochondrial | 0.81 | 7.1E-05 |
| A0A1S4C6F3_TOBAC | External alternative NAD(P)H-ubiquinone oxidoreductase B1, mitochondrial | 0.79 | 4.7E-04 |
| A0A1S4B9Y3_TOBAC | Uncharacterized protein At2g27730, mitochondrial-like | 0.62 | 4.6E-04 |
| A0A1S3X0W4_TOBAC | Cytochrome c1-2, heme protein, mitochondrial-like | 0.61 | 2.0E-03 |
| A0A1S3ZKM8_TOBAC | Ubiquinone biosynthesis protein COQ4 homolog, mitochondrial | 0.60 | 1.2E-02 |
| **Amino acid metabolism** | | | |
| SYE_TOBAC | Glutamate--tRNA ligase, chloroplastic/mitochondrial | -0.67 | 5.8E-05 |
| A0A1S3XY89_TOBAC | Arginase 1, mitochondrial | 2.01 | 7.5E-06 |
| A0A1S3Z2C6_TOBAC | Methylcrotonoyl-CoA carboxylase beta chain, mitochondrial-like | 1.71 | 2.5E-05 |

| | | | |
|---|---|---|---|
| A0A1S3ZD18_TOBAC | Isovaleryl-CoA dehydrogenase, mitochondrial | 1.63 | 7.1E-06 |
| A0A1S3XVN9_TOBAC | Mitochondrial amidoxime reducing component 2-like isoform X1 | 1.19 | 2.3E-04 |
| A0A1S3Z2K7_TOBAC | Methylcrotonoyl-CoA carboxylase subunit alpha, mitochondrial-like | 1.19 | 5.6E-05 |
| A0A1S4DR86_TOBAC | Hydroxymethylglutaryl-CoA lyase, mitochondrial-like | 1.04 | 6.0E-05 |
| A0A1S4C601_TOBAC | Alanine--glyoxylate aminotransferase 2 homolog 1, mitochondrial-like | 1.03 | 5.0E-05 |
| A0A1S4A1B1_TOBAC | Threonine--tRNA ligase, chloroplastic/mitochondrial 2 | 0.93 | 1.3E-04 |
| A0A1S4A4G8_TOBAC | Ferredoxin-dependent glutamate synthase 1, chloroplastic/mitochondrial-like | 0.89 | 4.5E-04 |
| A0A1S3Y512_TOBAC | Threonine--tRNA ligase, mitochondrial 1-like isoform X2 | 0.85 | 1.1E-02 |
| A0A1S3XS90_TOBAC | Bifunctional dethiobiotin synthetase/7,8-diamino-pelargonic acid aminotransferase, mitochondrial-like isoform X1 | 0.83 | 5.7E-04 |
| A0A1S4AZA4_TOBAC | Gamma aminobutyrate transaminase 1, mitochondrial isoform X1 | 0.80 | 7.6E-04 |
| A0A1S3ZGF8_TOBAC | Delta-1-pyrroline-5-carboxylate dehydrogenase 12A1, mitochondrial-like | 0.62 | 1.5E-04 |
| A0A1S3ZQY8_TOBAC | Bifunctional D-cysteine desulfhydrase/1-aminocyclopropane-1-carboxylate deaminase, mitochondrial | 0.61 | 5.5E-05 |
| **TCA cycle** | | | |
| A0A1S4BDU7_TOBAC | Acylpyruvase FAHD1, mitochondrial-like | 0.95 | 7.1E-05 |
| A0A1S4DJT3_TOBAC | Succinate dehydrogenase subunit 5, mitochondrial-like isoform X2 | 0.64 | 1.0E-03 |
| A0A1S4D199_TOBAC | Succinate dehydrogenase subunit 6, mitochondrial-like | 0.59 | 2.0E-03 |

*: t-test with FDR correction

**Table S10. DAPs involved in tubular ER network and root hair elongation**

| UniProtKB | Protein description | Log2 fold-change | *p*-value* |
|---|---|---|---|
| **Reticulons** | | | |
| A0A1S3XSN1_TOBAC | Reticulon-like protein | -1.77 | 5.7E-06 |
| A0A1S3XEV5_TOBAC | Rreticulon-like protein | -1.65 | 3.5E-05 |

| | | | |
|---|---|---|---|
| A0A1S4CBJ7_TOBAC | Reticulon-like protein | -1.36 | 1.6E-04 |
| **GTPases** | | | |
| A0A1S4BSP5_TOBAC | Nuclear/nucleolar GTPase 2 | -1.53 | 1.0E-04 |
| A0A1S4AAV4_TOBAC | LOW QUALITY PROTEIN: GTPase LSG1-2 | -1.04 | 1.1E-04 |
| A0A1S4DQ39_TOBAC | Protein ROOT HAIR DEFECTIVE 3 homolog | -0.82 | 4.9E-05 |
| A0A1S4BU20_TOBAC | Mitochondrial Rho GTPase | 0.70 | 2.0E-03 |
| **Actin** | | | |
| A0A1S3YSA4_TOBAC | Actin-like | -0.65 | 1.4E-04 |
| A0A1S3ZMD9_TOBAC | Actin | -0.62 | 1.4E-04 |
| **Actin binding proteins** | | | |
| A0A1S4BZF6_TOBAC | Villin-2-like | -1.27 | 5.3E-04 |
| A0A1S4DQI1_TOBAC | Formin-like protein 18 | -0.80 | 2.9E-04 |
| A0A1S4DAT8_TOBAC | Myosin-6-like | -0.65 | 5.9E-05 |
| A0A1S4BQ28_TOBAC | Myosin-1-like | -0.59 | 5.5E-04 |
| **Phospholipids** | | | |
| A0A1S4C3I1_TOBAC | Phosphatidylinositol 3,4,5-trisphosphate 3-phosphatase and protein-tyrosine-phosphatase PTEN2A-like | -1.32 | 1.2E-04 |
| A0A1S4CNE1_TOBAC | Phosphatidylinositol 4-kinase alpha 1 | -0.67 | 2.0E-03 |
| **Microtubule severing proteins** | | | |
| A0A1S4B058_TOBAC | Katanin p60 ATPase-containing subunit A-like 2 isoform X1 | -1.89 | 2.2E-04 |
| A0A1S4BXG6_TOBAC | Katanin p60 ATPase-containing subunit A1 | -1.03 | 2.0E-03 |
| **Calcium binding/dependent proteins** | | | |
| Q93YF4_TOBAC | Calcium-dependent protein kinase 2 | -1.27 | 1.5E-05 |
| A0A1S4CK71_TOBAC | Calcium load-activated calcium channel | -1.05 | 9.6E-05 |
| A0A1S4C2K7_TOBAC | Calcium-transporting ATPase | -1.00 | 5.9E-05 |
| A0A1S4BDN1_TOBAC | Calcium-transporting ATPase 4, endoplasmic reticulum-type-like | -0.82 | 2.8E-05 |
| A0A1S3ZRG7_TOBAC | Calcium-dependent protein kinase 5 | -0.78 | 2.9E-04 |
| A0A1S3XR50_TOBAC | Probable calcium-binding protein CML13 | -0.76 | 4.9E-04 |
| A0A1S3X4F2_TOBAC | Calcium-dependent protein kinase 4 isoform X2 | -0.70 | 8.2E-05 |
| A0A1S4BCT9_TOBAC | Calcium-dependent protein kinase 2-like | -0.69 | 1.9E-05 |
| A0A1S4AQV9_TOBAC | Probable calcium-binding protein CML14 | -0.62 | 4.6E-04 |
| A0A1S3YRP4_TOBAC | Probable calcium-binding protein CML49 | 1.82 | 5.1E-05 |
| A0A1S4DAE0_TOBAC | Probable calcium-binding protein CML50 | 0.88 | 6.0E-04 |
| **NADPH oxidase** | | | |
| Q8RVJ9_TOBAC | NADPH oxidase | -0.60 | 6.5E-04 |
| **Proton-ATPases** | | | |
| A0A1S3YUM1_TOBAC | V-type proton ATPase subunit a | -1.00 | 1.1E-04 |

| | | | |
|---|---|---|---|
| VATG2_TOBAC | V-type proton ATPase subunit G 2 | -0.96 | 1.3E-02 |
| **Vesicle-mediated transporters** | | | |
| A0A1S3XFQ2_TOBAC | Exocyst complex component EXO70A1-like | -0.87 | 7.5E-05 |
| A0A1S3XHT6_TOBAC | Protein transport protein sec16 | -1.00 | 2.8E-05 |
| A0A1S3YKI1_TOBAC | Protein transport protein SEC23-like | -0.65 | 1.9E-04 |
| A0A1S3YTI3_TOBAC | MAG2-interacting protein 2-like | -0.76 | 2.0E-03 |
| A0A1S3ZJ13_TOBAC | Clathrin light chain | -0.69 | 3.6E-05 |
| A0A1S3ZN00_TOBAC | Protein transport protein SEC23-like | -0.93 | 2.2E-04 |
| A0A1S3ZVR0_TOBAC | Rhomboid-like protein 19 | -1.16 | 3.4E-04 |
| A0A1S4A1T2_TOBAC | Protein transport protein SEC13 homolog B-like | -0.75 | 4.9E-04 |
| A0A1S4AU43_TOBAC | 25.3 kDa vesicle transport protein-like isoform X1 | -0.73 | 5.8E-05 |
| A0A1S4B101_TOBAC | TBC1 domain family member 2B-like isoform X1 | -0.70 | 2.1E-04 |
| A0A1S4B103_TOBAC | Auxilin-related protein 2-like | -0.60 | 9.0E-03 |
| A0A1S4B7H0_TOBAC | 25.3 kDa vesicle transport protein | -0.90 | 1.2E-04 |
| A0A1S4BHU2_TOBAC | Protein transport protein SEC24-like | -0.65 | 3.8E-04 |
| A0A1S4BS72_TOBAC | SEC1 family transport protein SLY1-like | -0.63 | 1.8E-04 |
| A0A1S4CVD2_TOBAC | Golgi SNAP receptor complex member 1 | -1.13 | 2.7E-04 |
| A0A1S4D9U5_TOBAC | Protein transport protein SEC23-like | -0.89 | 5.2E-04 |
| A0A1S4DH90_TOBAC | Coatomer subunit beta'-1-like | -0.64 | 8.6E-05 |
| A0A1S4DRM1_TOBAC | Protein transport protein Sec24-like At3g07100 | -0.76 | 4.3E-05 |
| Q9SDQ5_TOBAC | GTP-binding protein SAR1A-like | -0.67 | 4.3E-05 |

*: t-test with FDR correction

**Table S11. DAPs involved in ribosome biogenesis and ribosome assembly**

| UniProtKB | Protein description | Log2 fold-change | *p*-value* |
|---|---|---|---|
| **Ribosome biogenesis related proteins** | | | |
| A0A1S3Y752_TOBAC | Ribosome biogenesis protein NSA2 homolog | -1.59 | 2.5E-05 |
| A0A1S3ZMA0_TOBAC | Ribosome biogenesis protein BMS1 homolog | -1.31 | 1.4E-04 |
| A0A1S3Z5Y9_TOBAC | Ribosome biogenesis regulatory protein | -1.28 | 8.1E-05 |
| A0A1S4BK64_TOBAC | Ribosome biogenesis protein WDR12 homolog | -1.25 | 5.5E-05 |

| | | | |
|---|---|---|---|
| A0A1S3XGV3_TOBAC | Putative ribosome biogenesis protein C8F11.04 | -1.12 | 2.8E-05 |
| A0A1S3Z0I4_TOBAC | Ribosome biogenesis protein BOP1 homolog | -1.08 | 6.4E-05 |
| A0A1S4DAU9_TOBAC | Ribosome biogenesis protein BRX1 homolog | -0.76 | 3.7E-04 |
| **AAA-type ATPases** | | | |
| A0A1S4BN34_TOBAC | ATPase family AAA domain-containing protein 1-like | -1.26 | 1.0E-03 |
| A0A1S4CWQ5_TOBAC | AAA-ATPase ASD, mitochondrial-like | 0.92 | 3.0E-03 |
| A0A1S4BZ22_TOBAC | AAA-ATPase At3g28580-like | 0.72 | 9.0E-03 |
| **Helicases** | | | |
| A0A1S4DI29_TOBAC | DEAD-box ATP-dependent RNA helicase 10-like | -1.60 | 5.5E-05 |
| A0A1S3XE50_TOBAC | ATP-dependent RNA helicase-like protein DB10 isoform X1 | -1.49 | 1.0E-03 |
| A0A1S4A4Z1_TOBAC | DEAD-box ATP-dependent RNA helicase 9-like isoform X1 | -1.48 | 2.4E-05 |
| A0A1S4C7R7_TOBAC | DEAD-box ATP-dependent RNA helicase 7-like | -1.43 | 8.9E-05 |
| A0A1S3ZEG5_TOBAC | DEAD-box ATP-dependent RNA helicase 53-like isoform X1 | -1.40 | 3.4E-05 |
| A0A1S3YXI7_TOBAC | DEAD-box ATP-dependent RNA helicase 31-like | -1.33 | 1.6E-04 |
| A0A1S3YSR9_TOBAC | Putative DEAD-box ATP-dependent RNA helicase 29 | -1.31 | 6.0E-05 |
| A0A1S3XY80_TOBAC | ATP-dependent RNA helicase-like protein DB10 isoform X1 | -1.26 | 5.3E-05 |
| A0A1S4A617_TOBAC | ATP-dependent RNA helicase A-like protein | -1.22 | 1.2E-04 |
| A0A1S3XMH8_TOBAC | LOW QUALITY PROTEIN: DExH-box ATP-dependent RNA helicase DExH9-like | -1.12 | 1.4E-02 |
| A0A1S3YK03_TOBAC | DEAD-box ATP-dependent RNA helicase 13-like | -1.10 | 9.5E-04 |
| A0A1S3X981_TOBAC | DEAD-box ATP-dependent RNA helicase 57 isoform X1 | -0.97 | 4.1E-04 |
| A0A1S3YVT9_TOBAC | DEAD-box ATP-dependent RNA helicase 16-like | -0.77 | 3.0E-03 |
| A0A1S3ZS83_TOBAC | DEAD-box ATP-dependent RNA helicase 5-like | -0.72 | 1.6E-04 |
| **GTPases** | | | |
| A0A1S4BSP5_TOBAC | Nuclear/nucleolar GTPase 2 | -1.53 | 1.0E-04 |
| A0A1S3X678_TOBAC | Probable ADP-ribosylation factor GTPase-activating protein AGD6 | -1.41 | 5.9E-04 |

| | | | |
|---|---|---|---|
| A0A1S4AAV4_TOBAC | LOW QUALITY PROTEIN: GTPase LSG1-2 | -1.04 | 1.1E-04 |
| A0A1S4BU20_TOBAC | Mitochondrial Rho GTPase | 0.70 | 2.0E-03 |

*: t-test with FDR correction

**Table S12. DAPs involved in regulation of glutathione redox status**

| UniProtKB | Protein description | Log2 fold-change | *p*-value* |
|---|---|---|---|
| **Glutathione *S*-transferases** | | | |
| A0A1S3Y307_TOBAC | Glutathione S-transferase U10-like | 2.49 | 1.0E-03 |
| GSTX4_TOBAC | Probable glutathione S-transferase | 2.47 | 2.1E-04 |
| GSTX4_TOBAC | Probable glutathione S-transferase | 2.47 | 2.1E-04 |
| GSTX3_TOBAC | Probable glutathione S-transferase | 2.37 | 1.0E-03 |
| GSTXA_TOBAC | Probable glutathione S-transferase parA | 2.36 | 8.5E-04 |
| A0A1S4CQJ4_TOBAC | Glutathione S-transferase U9-like | 2.30 | 1.0E-03 |
| A0A1S4AJB2_TOBAC | Probable glutathione S-transferase | 2.23 | 1.0E-03 |
| A0A1S3YEE6_TOBAC | Glutathione S-transferase U9-like | 2.08 | 6.1E-04 |
| A0A1S3YUL8_TOBAC | Glutathione S-transferase-like | 2.02 | 9.1E-05 |
| A0A1S3X0J2_TOBAC | Glutathione S-transferase L3-like isoform X1 | 1.85 | 4.3E-05 |
| A0A1S4BDE5_TOBAC | Probable glutathione S-transferase | 1.76 | 1.0E-03 |
| GSTX1_TOBAC | Probable glutathione S-transferase | 1.73 | 5.3E-04 |
| GSTF2_TOBAC | Glutathione S-transferase APIC | 1.71 | 3.5E-04 |
| A0A1S4BZT5_TOBAC | Probable glutathione S-transferase | 1.70 | 1.0E-03 |
| Q40533_TOBAC | ParB product which have an activity of glutathione S-transferase | 1.66 | 5.8E-04 |
| A0A1S4AX14_TOBAC | Probable glutathione S-transferase | 1.64 | 1.0E-04 |
| A0A1S3YM66_TOBAC | Probable glutathione S-transferase parC | 1.64 | 3.0E-03 |
| A0A1S4BT19_TOBAC | Glutathione S-transferase zeta class-like | 1.55 | 2.0E-03 |
| A0A1S4BFF8_TOBAC | Probable glutathione S-transferase | 1.40 | 2.7E-04 |
| A0A1S3Z4V2_TOBAC | Glutathione S-transferase U8-like | 1.34 | 3.0E-03 |
| A0A1S4CJX9_TOBAC | Glutathione S-transferase T1-like | 1.32 | 1.0E-03 |
| A0A1S4CS55_TOBAC | Glutathione transferase GST 23-like | 1.31 | 3.0E-03 |
| A0A1S3XIX1_TOBAC | Probable glutathione S-transferase | 1.24 | 7.6E-04 |
| A0A1S3Y8V8_TOBAC | Glutathione S-transferase DHAR3, chloroplastic | 1.18 | 9.0E-03 |
| A0A1S4DMX8_TOBAC | Probable glutathione S-transferase | 0.88 | 9.0E-03 |
| A0A1S4A868_TOBAC | Glutathione S-transferase | 0.87 | 3.0E-03 |
| A0A1S4A502_TOBAC | Glutathione S-transferase T1-like | 0.85 | 3.9E-05 |

| | | | |
|---|---|---|---|
| A0A1S3YQ00_TOBAC | Microsomal glutathione S-transferase 3-like | 0.73 | 4.2E-04 |
| **Other glutathione metabolism related proteins** | | | |
| A0A1S3YKT8_TOBAC | Glutathione reductase, chloroplastic-like isoform X1 | 1.47 | 2.0E-04 |
| A0A1S4B721_TOBAC | Lactoylglutathione lyase | 1.06 | 4.1E-06 |
| A0A1S4AJD1_TOBAC | Probable lactoylglutathione lyase, chloroplastic isoform X1 | 0.90 | 6.0E-05 |
| A0A077D837_TOBAC | Glutathione synthetase | 0.71 | 8.8E-04 |
| A0A1S3XER8_TOBAC | S-formylglutathione hydrolase | 0.67 | 9.8E-05 |
| A0A1S3YB42_TOBAC | Glutathione peroxidase | 0.65 | 4.8E-04 |
| GSHRP_TOBAC | Glutathione reductase, chloroplastic | 0.62 | 7.6E-05 |
| A0A1S3XF65_TOBAC | Lactoylglutathione lyase | 0.61 | 1.4E-04 |

*: t-test with FDR correction

**Table S13. RNA polymerases involved in transcription**

| UniProtKB | Protein description | Log2 fold-change | *p*-value* |
|---|---|---|---|
| A0A1S4BU78_TOBAC | RNA polymerase II-associated protein 3-like | -1.11 | 1.4E-04 |
| A0A1S4CS69_TOBAC | DNA-directed RNA polymerases IV and V subunit 4-like | -0.73 | 5.0E-04 |
| A0A1S3XKG9_TOBAC | DNA-directed RNA polymerase I subunit rpa49-like isoform X1 | -0.64 | 1.0E-03 |
| A0A1S4DES6_TOBAC | DNA-directed RNA polymerases I and III subunit RPAC2-like isoform X1 | -0.62 | 2.5E-04 |
| A0A1S4BBD8_TOBAC | DNA-directed RNA polymerase I subunit 1-like | -0.61 | 2.0E-03 |
| A0A1S3YG40_TOBAC | DNA-directed RNA polymerases I and III subunit rpac1-like isoform X2 | -0.60 | 1.6E-04 |

*: t-test with FDR correction

**Table S14. Serine/threonine-protein kinases and phosphatases involved in cell proliferation and expansion**

| UniProtKB | Protein Description | Log2 fold-change | *p*-value* |
|---|---|---|---|
| A0A1S4A212_TOBAC | Serine/threonine-protein kinase 38-like | -1.72 | 1.0E-03 |

| | | | |
|---|---|---|---|
| A0A1S4AK92_TOBAC | Serine/threonine-protein kinase SAPK7-like | -1.39 | 5.6E-05 |
| A0A1S3YRK2_TOBAC | Probable LRR receptor-like serine/threonine-protein kinase At1g05700 | -1.33 | 4.6E-04 |
| A0A1S3ZMS6_TOBAC | Serine/threonine-protein kinase 16-like | -1.25 | 4.9E-04 |
| A0A1S4BJC6_TOBAC | Serine/threonine-protein kinase RUNKEL | -1.11 | 1.4E-02 |
| A0A1S3YND7_TOBAC | Probable LRR receptor-like serine/threonine-protein kinase At4g20940 isoform X1 | -1.08 | 1.0E-03 |
| A0A1S4DKM3_TOBAC | Serine/threonine-protein kinase HT1-like | -1.01 | 2.0E-03 |
| G5ELZ4_TOBAC | Serine/threonine-protein phosphatase 5 | -0.83 | 1.1E-04 |
| A0A1S3ZN95_TOBAC | Putative leucine-rich repeat receptor-like serine/threonine-protein kinase At2g14440 | -0.82 | 3.2E-04 |
| A0A1S3ZLP5_TOBAC | Probable serine/threonine-protein kinase DDB_G0280111 | -0.81 | 2.0E-03 |
| A0A1S4D0Z6_TOBAC | Serine/threonine-protein kinase EDR1-like | 0.76 | 2.0E-03 |

*: t-test with FDR correction

**Table S15. DAPs involved in folate and one carbon metabolisms**

| UniProtKB | Protein description | Log2 fold-change | *p*-value* |
|---|---|---|---|
| **Folate and one carbon metabolism** | | | |
| A0A1S3WYB9_TOBAC | GTP cyclohydrolase 1 | -2.64 | 2.0E-03 |
| A0A1S3Y8H2_TOBAC | Probable methyltransferase PMT8 | -1.70 | 3.0E-03 |
| A0A1S4AR74_TOBAC | Probable methyltransferase PMT20 | -1.70 | 3.1E-04 |
| A0A1S4B3D7_TOBAC | Uncharacterized protein At3g49720-like | -1.64 | 1.1E-04 |
| A0A1S4BH87_TOBAC | Probable methyltransferase PMT5 isoform X1 | -1.61 | 3.3E-05 |
| A0A1S3XT44_TOBAC | rRNA adenine N(6)-methyltransferase | -1.59 | 4.1E-04 |
| A0A1S3YJW9_TOBAC | Histone-lysine N-methyltransferase, H3 lysine-9 specific SUVH4-like | -1.41 | 4.1E-04 |
| A0A1S3ZLI4_TOBAC | Probable methyltransferase PMT13 | -1.41 | 5.4E-05 |
| A0A1S4BNH6_TOBAC | Probable methyltransferase PMT2 | -1.40 | 1.3E-04 |
| A0A1S3Y8F0_TOBAC | Probable methyltransferase PMT1 | -1.37 | 7.8E-05 |
| A0A1S3WXP6_TOBAC | Probable methyltransferase PMT3 | -1.35 | 2.2E-04 |
| A0A1S3ZFC4_TOBAC | Probable methyltransferase PMT26 | -1.33 | 1.7E-04 |
| W8TFR1_TOBAC | GTP cyclohydrolase II | -1.32 | 2.5E-05 |
| A0A1S4D663_TOBAC | Probable methyltransferase PMT14 | -1.32 | 4.2E-05 |
| A0A1S4CPT8_TOBAC | Probable 18S rRNA (Guanine-N(7))-methyltransferase | -1.17 | 5.9E-04 |
| A0A1S3XBX4_TOBAC | Phosphoethanolamine N-methyltransferase 1-like | -1.16 | 3.1E-05 |

| | | | |
|---|---|---|---|
| A0A1S4A3W8_TOBAC | (RS)-norcoclaurine 6-O-methyltransferase-like | -1.05 | 5.0E-03 |
| A0A1S3XLU7_TOBAC | tRNA (Cytosine(34)-C(5))-methyltransferase-like | -1.04 | 4.6E-04 |
| A0A1S3X806_TOBAC | Putative rRNA methyltransferase | -1.01 | 2.4E-04 |
| A0A1S3ZBS6_TOBAC | Methyltransferase-like protein 13 | -0.96 | 8.1E-05 |
| A0A1S4CZU1_TOBAC | Probable protein arginine N-methyltransferase 1 | -0.94 | 3.8E-05 |
| A0A1S4BY02_TOBAC | Probable protein arginine N-methyltransferase 3 | -0.90 | 3.5E-04 |
| A0A1S4B9Y5_TOBAC | Alpha N-terminal protein methyltransferase 1-like isoform X1 | -0.89 | 4.2E-04 |
| A0A1S3Y6U4_TOBAC | Glucuronoxylan 4-O-methyltransferase 3-like | -0.89 | 5.1E-04 |
| A0A1S4CCM9_TOBAC | Protein arginine N-methyltransferase | -0.88 | 4.8E-05 |
| A0A1S3Y1B2_TOBAC | Probable pectin methyltransferase QUA2 | -0.86 | 7.7E-04 |
| A0A1S3XBQ8_TOBAC | Probable 28S rRNA (Cytosine-C(5))-methyltransferase | -0.82 | 4.6E-04 |
| A0A1S3X4I9_TOBAC | 2-Methyl-6-phytyl-1,4-hydroquinone methyltransferase, chloroplastic-like | -0.75 | 2.5E-05 |
| A0A1S4CW49_TOBAC | Probable methyltransferase PMT15 | -0.75 | 1.7E-04 |
| CAMT2_TOBAC | Caffeoyl-CoA O-methyltransferase 2 | -0.74 | 4.0E-03 |
| A0A1S4ASH5_TOBAC | Protein arginine N-methyltransferase PRMT10-like | -0.73 | 2.7E-04 |
| A0A1S3ZEV6_TOBAC | tRNA (Guanine(10)-N2)-methyltransferase homolog | -0.69 | 3.0E-03 |
| A0A1S4C1X0_TOBAC | tRNA (guanine-N(7)-)-methyltransferase | -0.65 | 5.5E-04 |
| A0A1S4DN28_TOBAC | DNA (Cytosine-5)-methyltransferase DRM2-like isoform X1 | -0.64 | 3.0E-03 |
| A0A1S3X4H4_TOBAC | tRNA (guanine(26)-N(2))-dimethyltransferase | -0.63 | 3.9E-04 |
| A0A1S3YCG8_TOBAC | Putative methyltransferase NSUN6 isoform X2 | -0.61 | 2.0E-03 |
| A0A1S4B0J2_TOBAC | Probable methyltransferase PMT11 | -0.60 | 3.0E-03 |
| PMT4_TOBAC | Putrescine *N*-methyltransferase 4 | -0.60 | 1.5E-02 |
| A0A1S3YAK7_TOBAC | Probable methyltransferase PMT28 | -0.59 | 5.0E-03 |
| A0A1S3YGI7_TOBAC | Putative methyltransferase C9orf114 isoform X1 | -0.59 | 1.9E-04 |
| A0A1S4D868_TOBAC | Putative methyltransferase DDB_G0268948 | 3.02 | 9.9E-05 |
| A0A1S4A958_TOBAC | Probable caffeoyl-CoA O-methyltransferase At4g26220 | 1.01 | 1.0E-03 |
| A0A1S3YDV3_TOBAC | Methionine S-methyltransferase | 0.73 | 1.3E-05 |

*: t-test with FDR correction

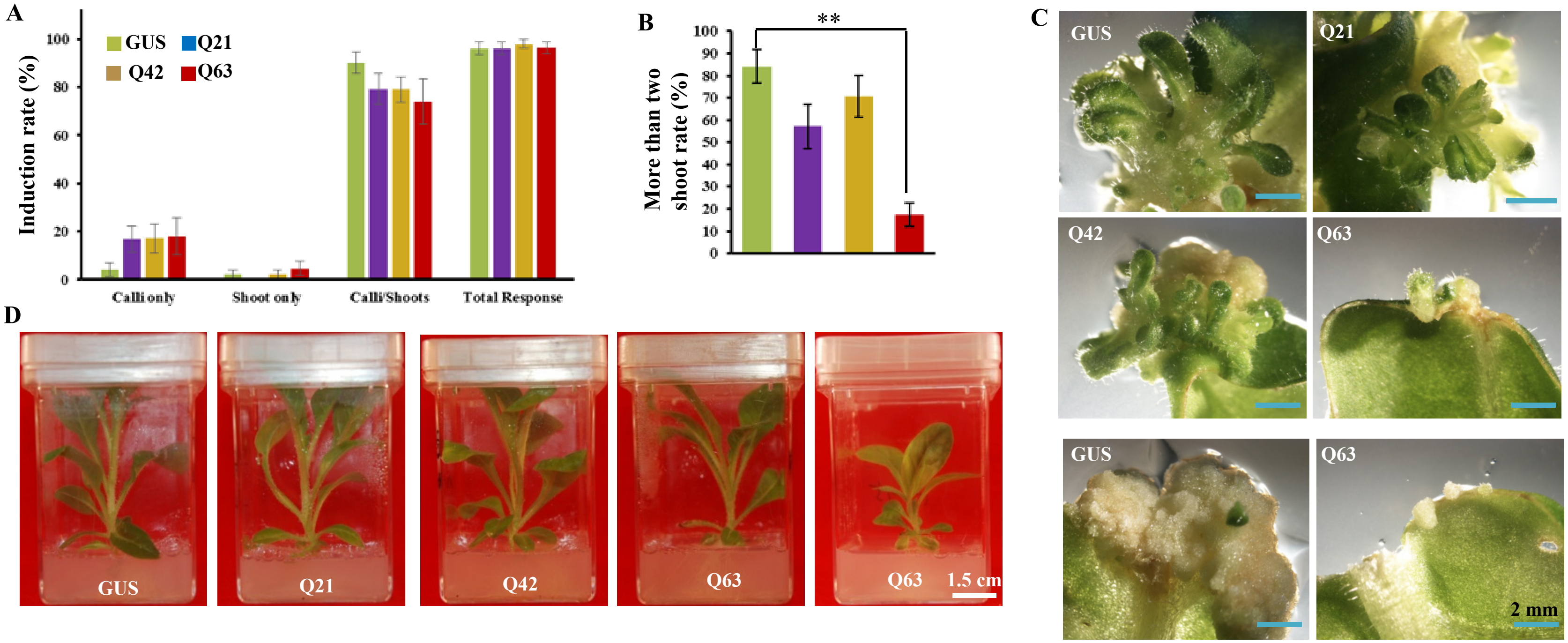


**Fig. S1. Transformation efficiency and regenerated transgenic plants. (A)** Culture response was counted as the percentage of regenerated calli, shoots, or both from each explant at week four of post-inoculation onto kanamycin selection medium. The average frequency of four plates (five explants per plate) from each genetic cassette per experiment was calculated. Data plotted was the average of three independent experiments ± SD. **(B)** The frequency of explants bearing more than two shoots per shoot formation site was calculated. Data plotted was the average of three independent experiments ± SD. **: significant differences at $p < 0.01$ level. **(C)** Induced shoots from four genetic cassettes, and induced calli from GUS compared to those from $Htt_{ex1}$Q63. **(D)** Representatives of kanamycin-resistant plants from four genetic cassettes were grown on maintenance medium under ~60 µmol $m^{-2}$ $s^{-1}$ light intensity. Plants carrying GUS, $Htt_{ex1}$Q21, $Htt_{ex1}$Q42 grew normally while plants carrying $Htt_{ex1}$Q63 showed some growth defects. Q21: $Htt_{ex1}$Q21; Q42: $Htt_{ex1}$Q42; and Q63: $Htt_{ex1}$Q63.

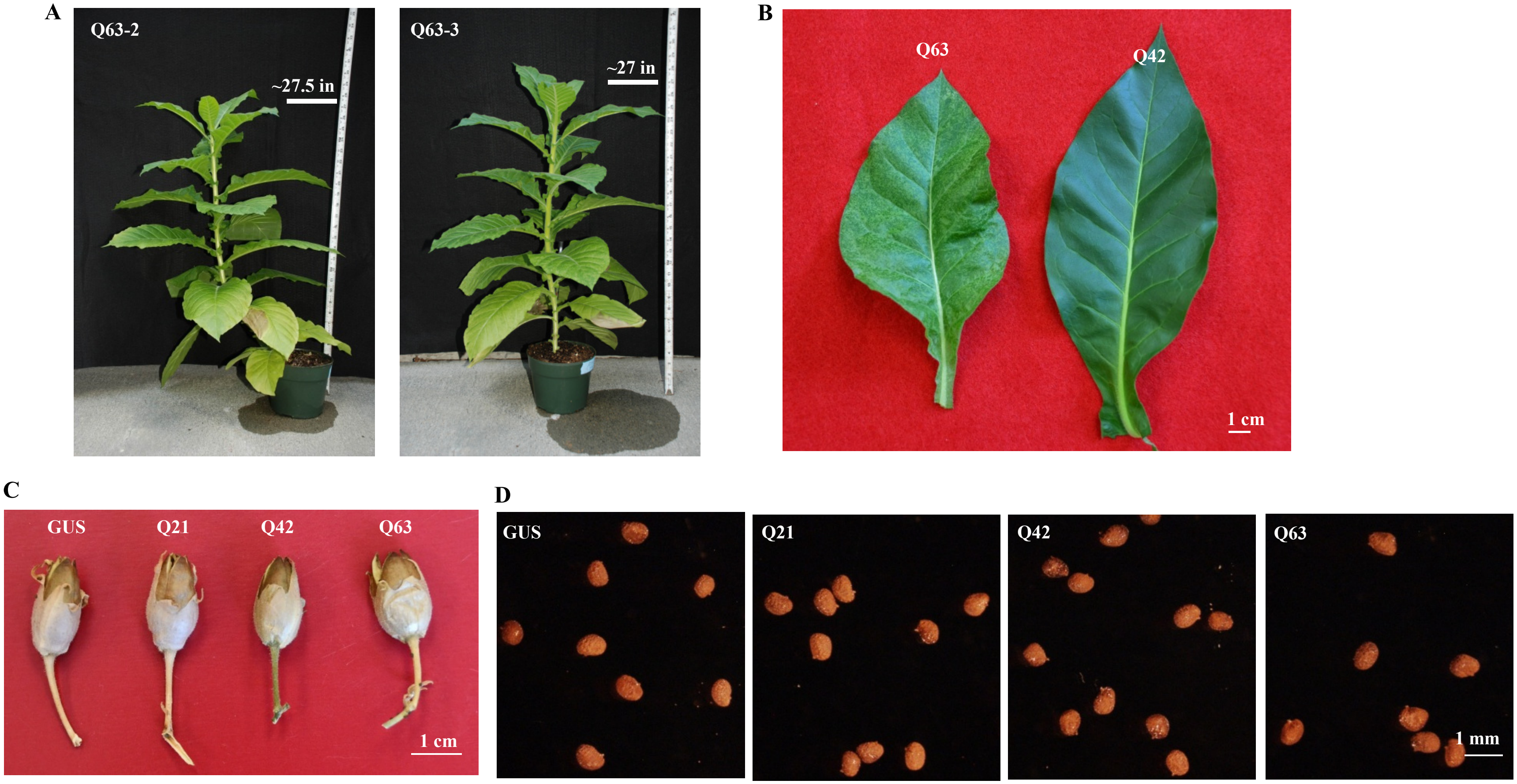


**Fig. S2. Transgenic plants, leaves, seed pods and seeds. (A)** Htt$_{ex1}$Q63 transgenic plants grew in soil under conditions of ~15 µmol m$^{-2}$ s$^{-1}$ photo-period (16 h d$^{-1}$) at 23°C. After eight weeks, median plant height reached approximately 20 to 30 inches (in). **(B)** Representatives of young leaves from soil grown Htt$_{ex1}$Q63 and Htt$_{ex1}$Q42 transgenic plants under conditions of ~15 µmol m$^{-2}$ s$^{-1}$ photo-period (16 h d$^{-1}$) at 23°C. Color defective leaf was observed only from Htt$_{ex1}$Q63 but not Htt$_{ex1}$Q42 plants. **(C)** Seed pods of GUS, Htt$_{ex1}$Q21, Htt$_{ex1}$Q42 and Htt$_{ex1}$Q63 transgenic lines were similar in sizes. **(D)** Seeds from the four transgenic seed pods shown in C were also similar in size and shape. Q21: Htt$_{ex1}$Q21; Q42: Htt$_{ex1}$Q42; and Q63: Htt$_{ex1}$Q63.

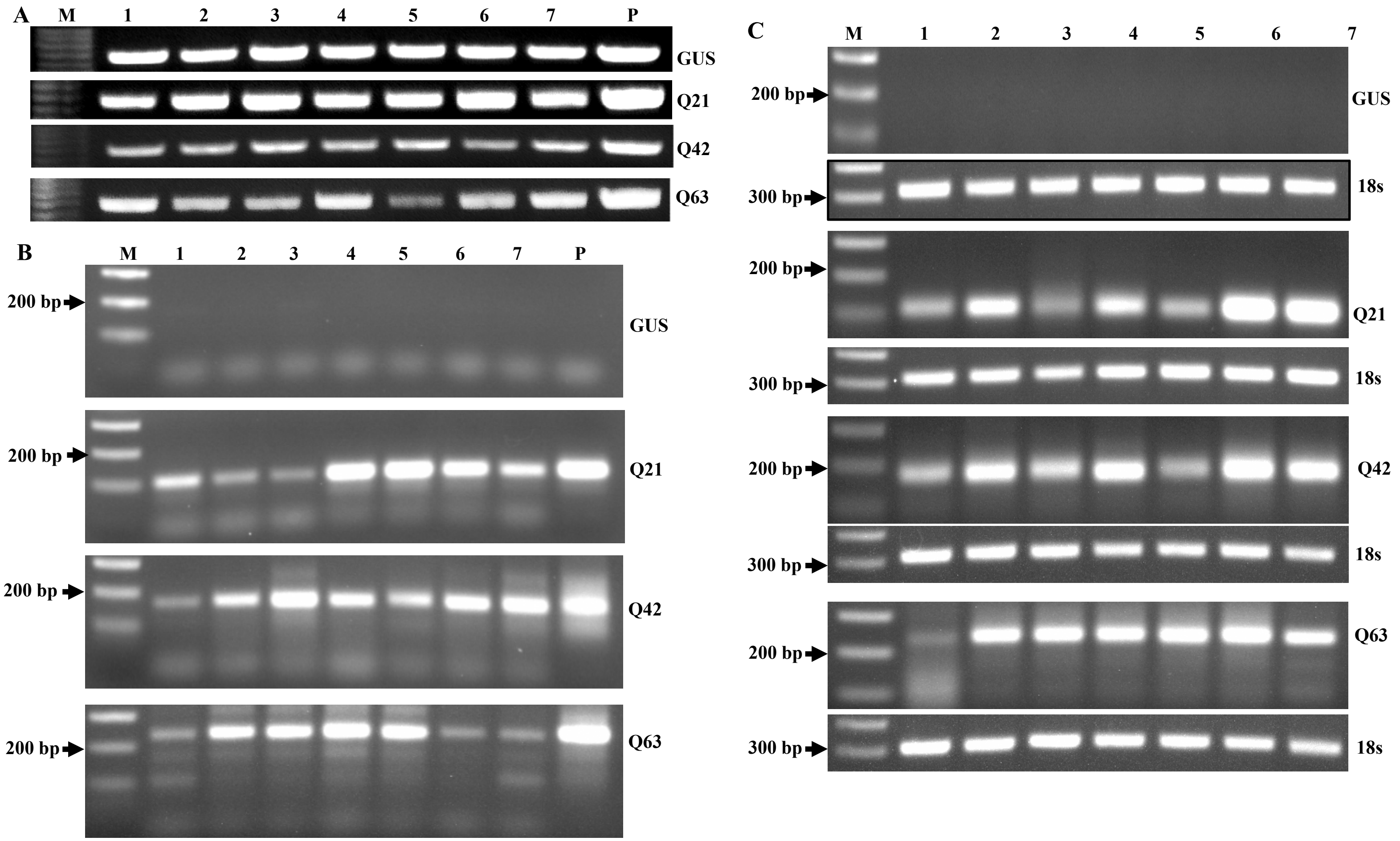


**Fig. S3. PCR and RT-PCR results of seven transgenic lines per genetic cassette. (A)** Genomic PCR of *nptII* performed with primers NPT-II 5'/NPT-II 3'. All had 795 bp PCR products. **(B)** Genomic PCR of *Htt* exon 1 with different "CAG" repeats performed with primers HDJamaF/HDJamaR2. $Htt_{ex1}Q21$, $Htt_{ex1}Q42$ and $Htt_{ex1}Q63$ transgenic lines had ~110 bp, ~170 bp and ~240 bp PCR products, respectively. GUS lines were negative of PCR. **(C)** RT-PCR of *Htt* exon 1 transcripts with different "CAG" repeats performed using primers HDJamaF/HDJamaR2 whose amplification covered the "CAG" repeat region. *18S* rRNA was used to show equal cDNA used for RT-PCR. $Htt_{ex1}Q21$, $Htt_{ex1}Q42$ and $Htt_{ex1}Q63$ transgenic lines had ~110 bp, ~170 bp and ~240 bp PCR products, respectively. GUS transgenic lines were negative without PCR products. P: Plasmid DNA. M: DNA ladder. Q21: $Htt_{ex1}Q21$; Q42: $Htt_{ex1}Q42$; and Q63: $Htt_{ex1}Q63$.

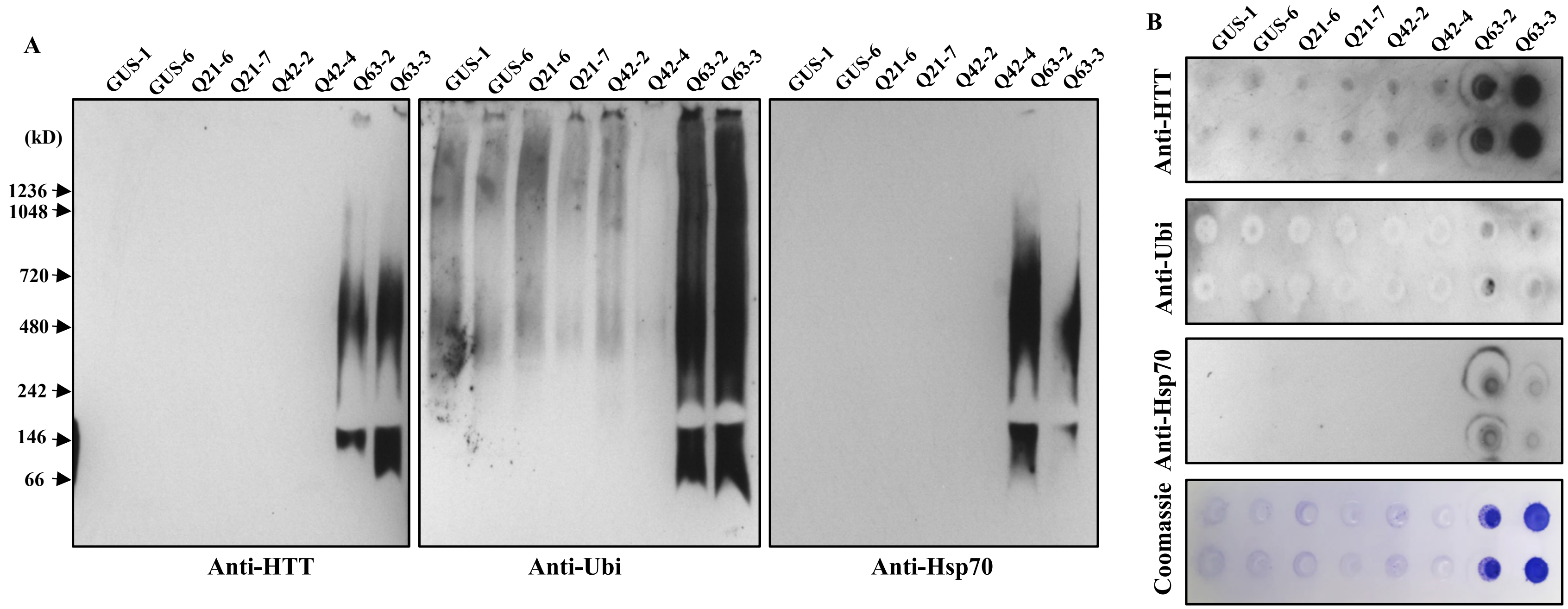


**Fig. S4. Detection of protein aggregates in plant leaves. (A)** Immunoblotting analysis of protein aggregates in young leave from two lines per genetic cassette by BN-PAGE. **(B)** Filter retardation assay of protein aggregates trapped on 0.2 µm membranes. One membrane was stained with coomassie R-250 (Coomassie) showing the presence of protein aggregates. Anti-HTT: anti-Huntingtin; Anti-Ubi: anti-Ubiquitin; and Anti-Hsp70: anti-Hsp70-biotin. Q21: $Htt_{ex1}$Q21; Q42: $Htt_{ex1}$Q42; and Q63: $Htt_{ex1}$Q63.

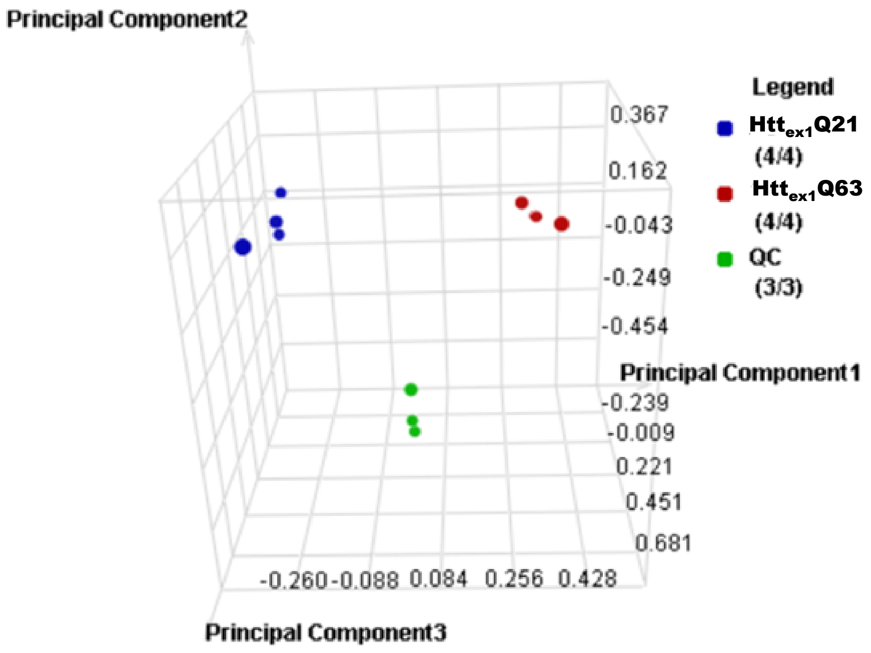


**Fig. S5. Principal component analysis of quantified proteins.** The expression values of the 5,073 proteins quantified by 2 or more peptides were Z-score-normalized followed by PCA using Rosetta Elucidator.

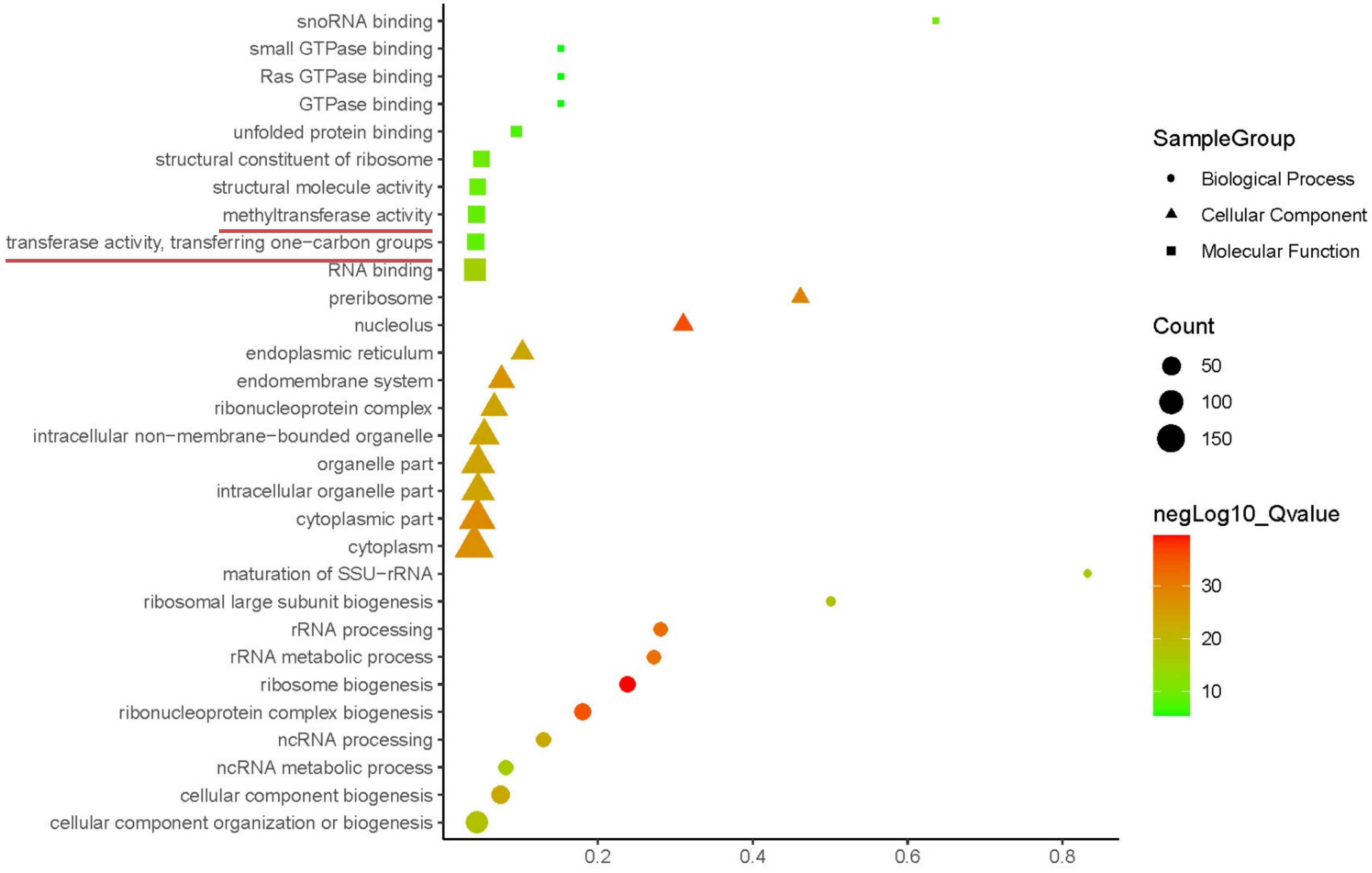


**Fig. S6. Selected top 10 GO results of abundance-decreased DAPs**.

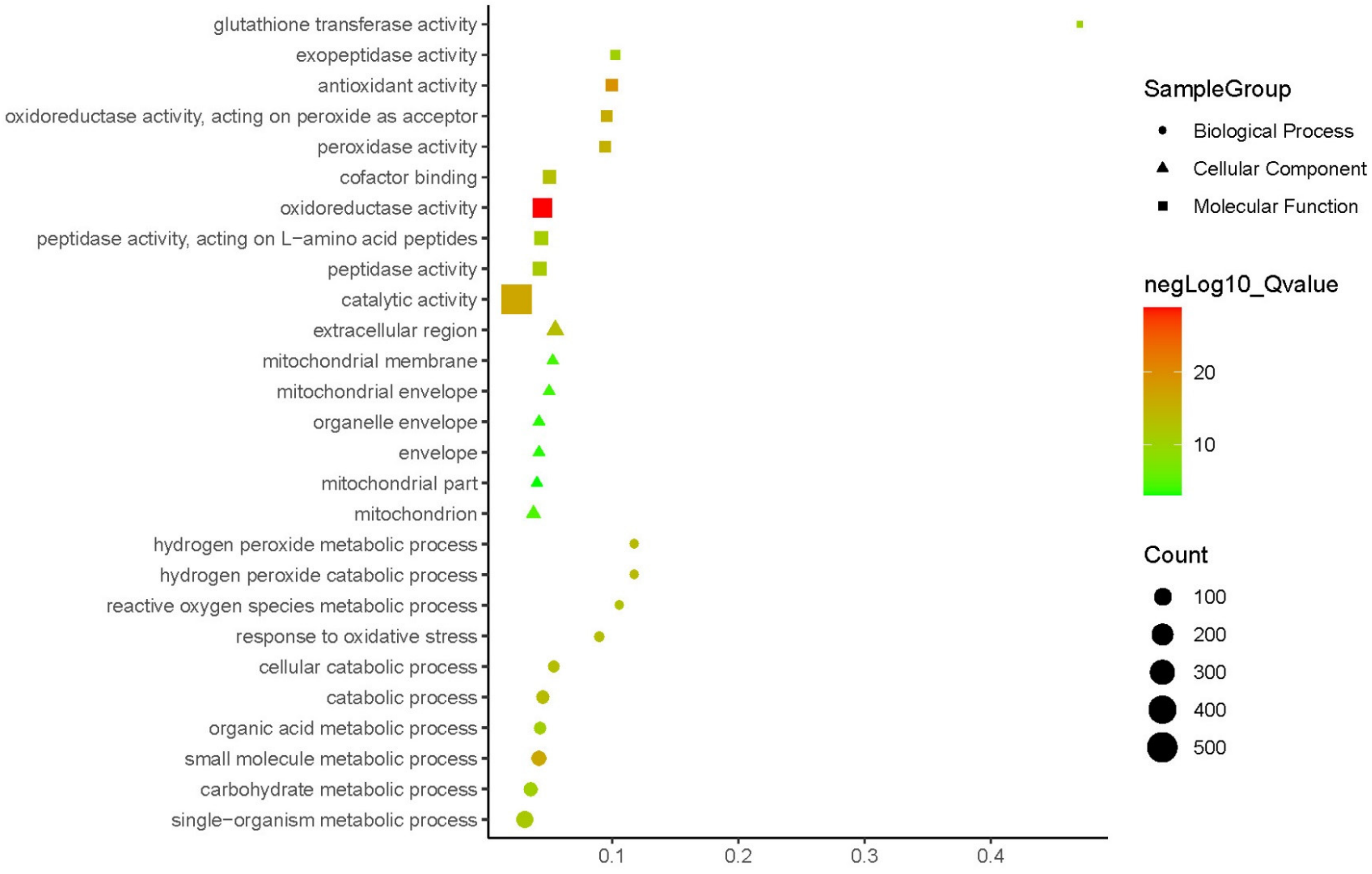


**Fig. S7. Selected top 10 GO results of GO results of abundance-increased DAPs**. For cellular component category, there are only 7 GO terms.

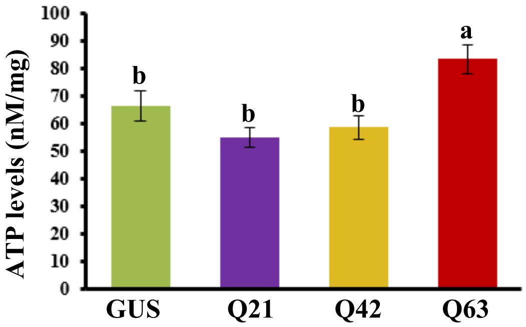


**Fig. S8. ATP levels of young leaf tissues.** ATP levels of young leaf tissues from four types of transgenic plants. Bioluminescent Assay Kit was used. Data plotted are the average of five biological repeats ± SD (n = 5). Different letters represent significant differences at $p < 0.05$ level.

A

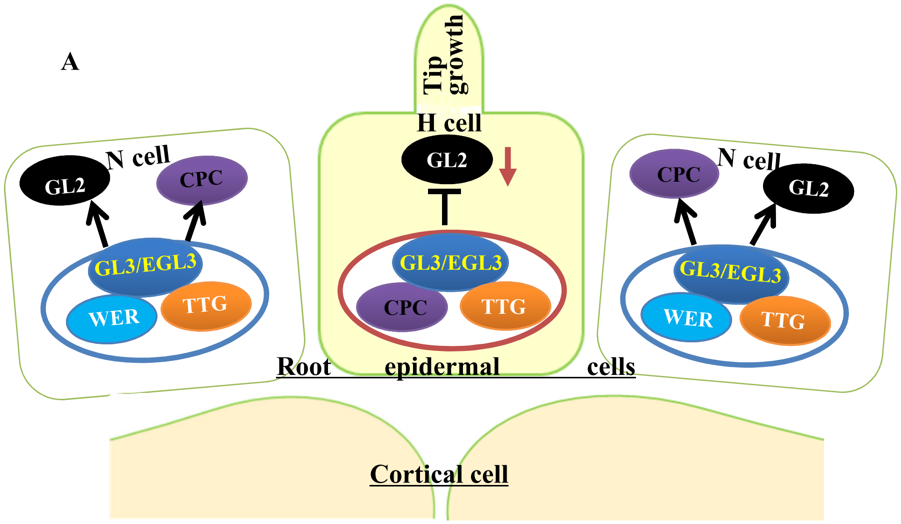


B

GL2 (Q63 vs. Q21 roots): -1.51-fold

C

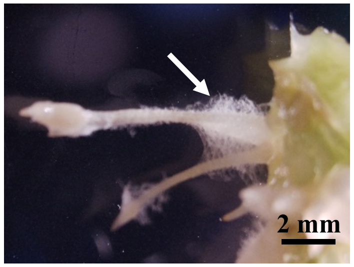


**Fig. S9. Root hair initiation in induced roots. (A)** Schematic diagram showing the differentiation of epidermal cells with related transcription factors adopted from Libault et al.[30]. The root epidermal cell can be differentiated into a root hair (H) or non-hair (N) cell. The process is determined by a group of transcription factors: GLABRA2 (GL2), CAPRICE (CPC), TRANSPARENT TESTA GLABRA (TTG), GLABRA3 (GL3)/ENHANCER OF GLABRA3 (EGL3) and WEREWOLF (WER). An epidermal cell becomes a H cell when GL2 was inhibited by the formation of CPC-TTG-GL3-EGL3 complex while it becomes a N cell when GL2 was induced by the formation of WER-TTG-GL3-EGL3. The red arrow indicates GL2 abundance was decreased. **(B)** GL2 protein level in $Htt_{ex1}$Q63 roots was reduced of 1.5-fold compared to $Htt_{ex1}$Q21 roots. **(C)** Induced roots having hairs when shoots were inoculated on the surface of medium.

A

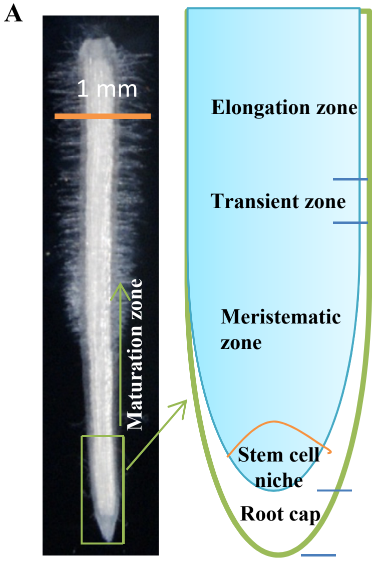


B

| Accession # | Protein description | FC* | Function** |
|---|---|---|---|
| Q9AYN9 | NQK1 MAPKK | -3.2 | Cell division[82] |
| A0A1S4C0N4 | Cyclin-dependent kinases regulatory subunit | -3 | Cell division[84] |
| A0A1S4B058 | Katanin p60 ATPase-containing subunit A-like 2 isoform X1 | -3.7 | Cell elongation[79] |
| A0A1S4BXG6 | Katanin p60 ATPase-containing subunit A1 | -2 | Cell elongation[79] |
| Q8GZV0 | Obtusifoliol-14-demethylase | -3.4 | Cell elongation[81] |
| A0A1S3X0S9 | Probable beta-1,4-xylosyltransferase IRX10L | -2.8 | Cell elongation[83] |
| A0A1S3YHB2 | Wall-associated receptor kinase 2-like | -2.3 | Cell expansion[80] |
| A0A1S4CA37 | Chitinase-like protein 1 | -3.4 | Cell expansion[84] |
| A0A1S3YBC1 | Protein ROOT INITIATION DEFECTIVE 3-like | -1.9 | Cell expansion[34] |

C

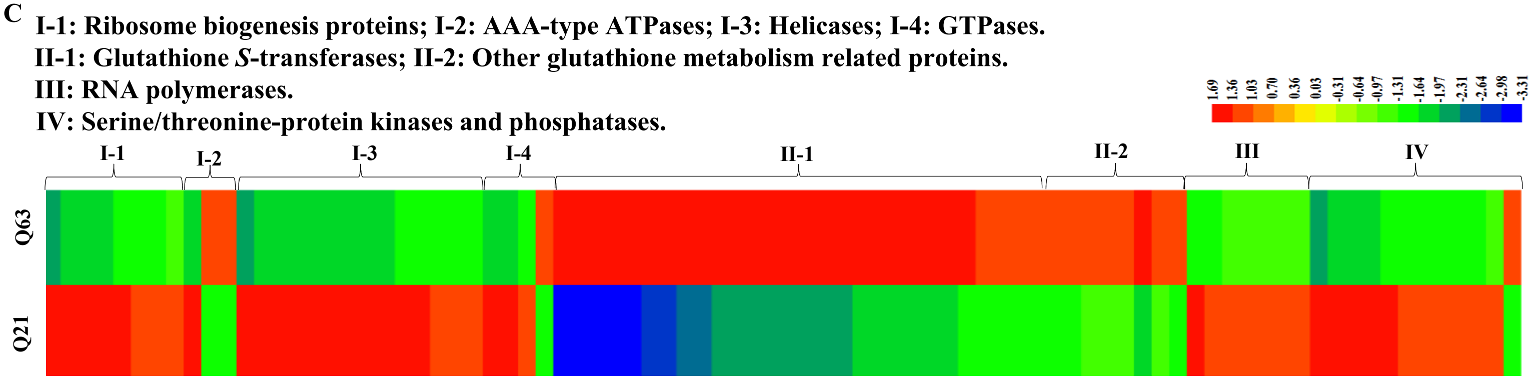


**Fig. S10. Normal root tip structure and DAPs associated with root cell division, elongation or expansion. (A)** Normal root tip structure. **(B)** DAPs associated with root cell division, elongation or expansion. *: Fold changes of $Htt_{ex1}$Q63 (Q63) vs. $Htt_{ex1}$Q21 (Q21); **: the references cited. **(C)** Four groups (I - IV) of DAPs associated with root growth and root cap integrity. Expression values are shown as a color scale at the right corner of the figure.

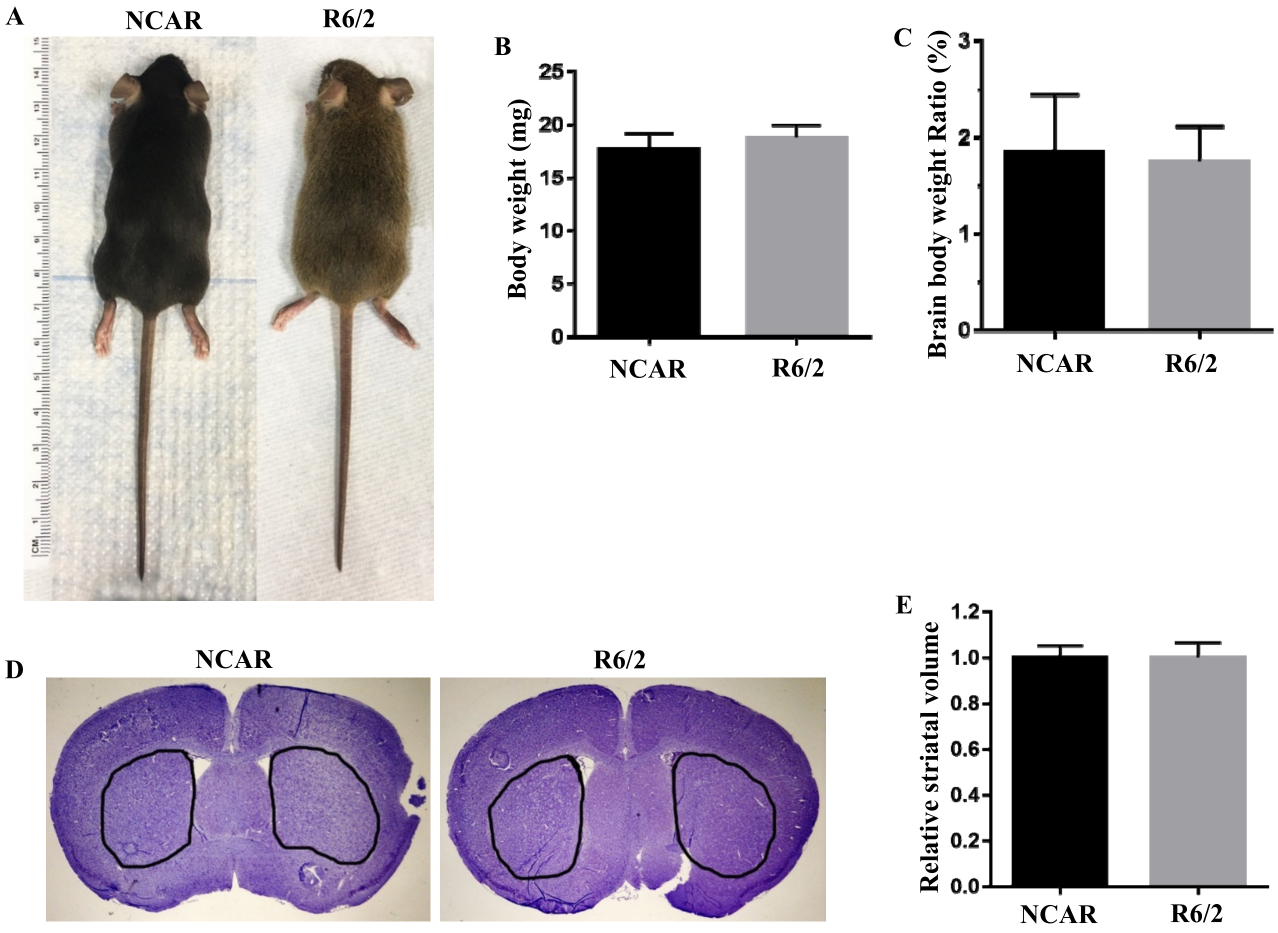


**Fig. S11. Characterization of 4-week old wide-type non-carrier (NCAR) and R6/2 HD mice. (A)** Mouse phenotype. **(B)** Body weight (n = 7). **(C)** Brain body weight ratio (n = 7). **(D)** Brain section with striatum region circled. **(E)** Relative striatal volume.

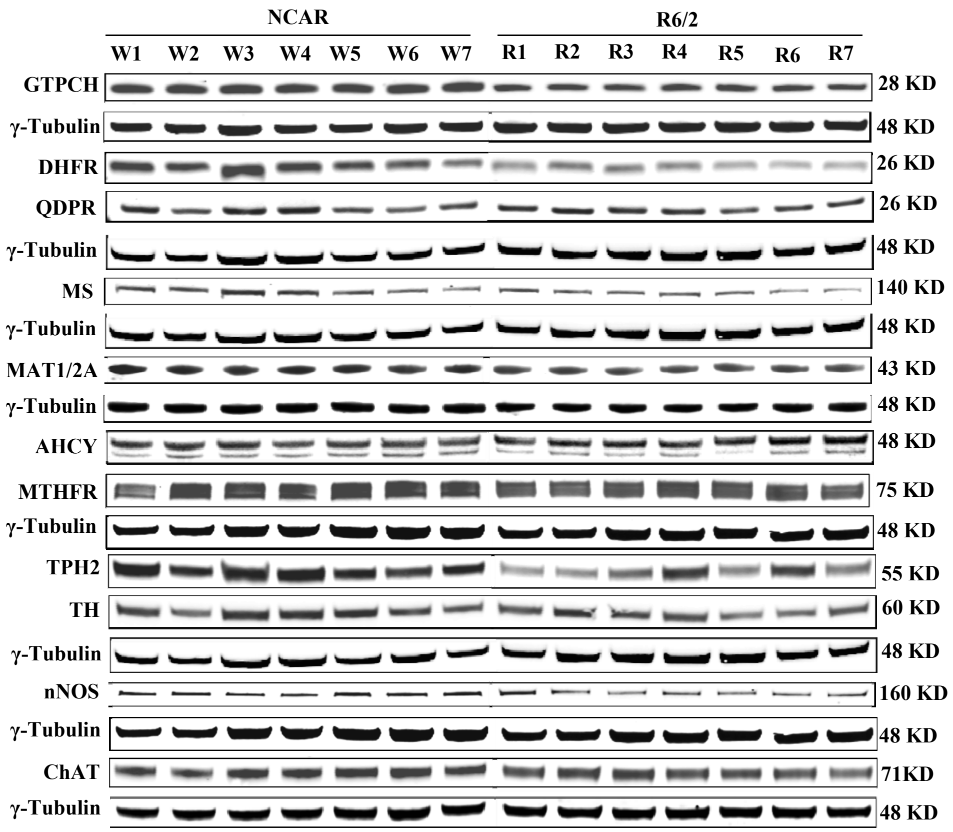


**Fig. S12. Representative Western blotting results of GTPCH, DHFR, QDPR, MS, MAT1/2A, AHCY, MTHFR, TPH2, TH, nNOS and ChAT.** Four-week-old male R6/2 and NCAR mice were used (n = 7). γ-Tubulin was used as internal control. Abbreviation used for enzymes: AHCY, *S*-adenosylhomocystein hydrolase; ChAT, choline acetyltransferase; DHFR, Dihydrofolate reductase; GTPCH, GTP cyclohydrolase I; MAT1/2A, methionine adenosyltransferase; MS, methionine synthase; MTHFR, methylene-tetrahydrofolate reductase; nNOS, neuronal nitric oxide synthase; QDPR, Quinoid dihydropteridine reductase; TH, tyrosine hydroxylase (Tyr); TPH2, tryptophan hydroxylase.
